# The barley mutant *multiflorus2.b* reveals quantitative genetic variation for new spikelet architecture

**DOI:** 10.1101/2021.04.06.438555

**Authors:** Ravi Koppolu, Guojing Jiang, Sara G Milner, Quddoos H Muqaddasi, Twan Rutten, Axel Himmelbach, Nils Stein, Martin Mascher, Thorsten Schnurbusch

## Abstract

Understanding the genetic basis of yield forming factors in small grain cereals is of extreme importance, especially in the wake of stagnation of further yield gains in these crops. One such yield forming factor in these cereals is the number of grain-bearing florets produced per spikelet. Wildtype barley (*Hordeum vulgare* L.) spikelets are determinate structures, the spikelet axis (rachilla) degenerates after producing single floret. In contrast, the rachilla of wheat (*Triticum ssp*.) spikelets, which are indeterminate, elongates to produce up to 12 florets. In our study, we characterized the barley spikelet determinacy mutant *multiflorus2.b* (*mul2.b*) that produced up to three fertile florets on elongated rachillae of lateral spikelets. Apart from the lateral spikelet indeterminacy (LS-IN), we also characterized the supernumerary spikelet phenotype in the central spikelets (CS-SS) of *mul2.b*. Through our phenotypic and genetic analyses, we identified two major QTLs on chromosomes 2H and 6H, and two minor QTLs on 3H for the LS-IN phenotype. For, the CS-SS phenotype we identified one major QTL on 6H, and a minor QTL on 5H chromosomes. Notably, the 6H QTLs for CS-SS and LS-IN phenotypes co-located with each other, potentially indicating that a single genetic factor might regulate both phenotypes. Thus, our in-depth phenotyping combined with genetic analyses revealed the quantitative nature of the LS-IN and CS-SS phenotypes in *mul2.b*, paving the way for cloning the genes underlying these QTLs in the future.

**Key message:** Spikelet indeterminacy and supernumerary spikelet phenotypes in barley *multiflorus2.b* mutant show polygenic inheritance. Genetic analysis of *multiflorus2.b* revealed major QTLs for spikelet determinacy and supernumerary spikelet phenotypes on 2H and 6H chromosomes.

## Introduction

Grass inflorescences are constituted by the ordered arrangement of spikelets (specialized flower-bearing structures) either directly on the inflorescence axis (rachis) or on the branches differentiated from the inflorescence axis. The Triticeae grasses display characteristic spike-shaped inflorescences with sessile spikelets attached to the inflorescence axis in a distichous phyllotaxis (Bonnett 1935, 1936; Kellogg et al. 2013). Among *Triticeae*, wheat (*Triticum spp*.) and barley (*Hordeum spp*.) represent the most economically important temperate cereal species (Schnurbusch 2019). Based on the architectural differences at the spikelet and spike levels, wheat and barley inflorescences can be classified into (i) determinate or (ii) indeterminate spikes (Bonnett 1966). In the determinate spikes of wheat, the inflorescence apex culminates with a terminal spikelet, producing a fixed number of spikelets per inflorescence. However, the spikelets in the wheat spike are indeterminate with multiple flowers (florets; up to 12), each produced at the articulation junction of the spikelet axis called rachilla (**Fig. 1E)**. In contrast, the inflorescence apex of barley remains indeterminate (no terminal spikelet) and the rachilla ceases to elongate upon initiation of the first floret and remains as a suppressed structure rendering determinacy to the spikelet (**Fig. 1B**). Thus, the spikelet determinacy or indeterminacy is dependent upon the extent of rachilla suppression or elongation.

**Figure 1.**
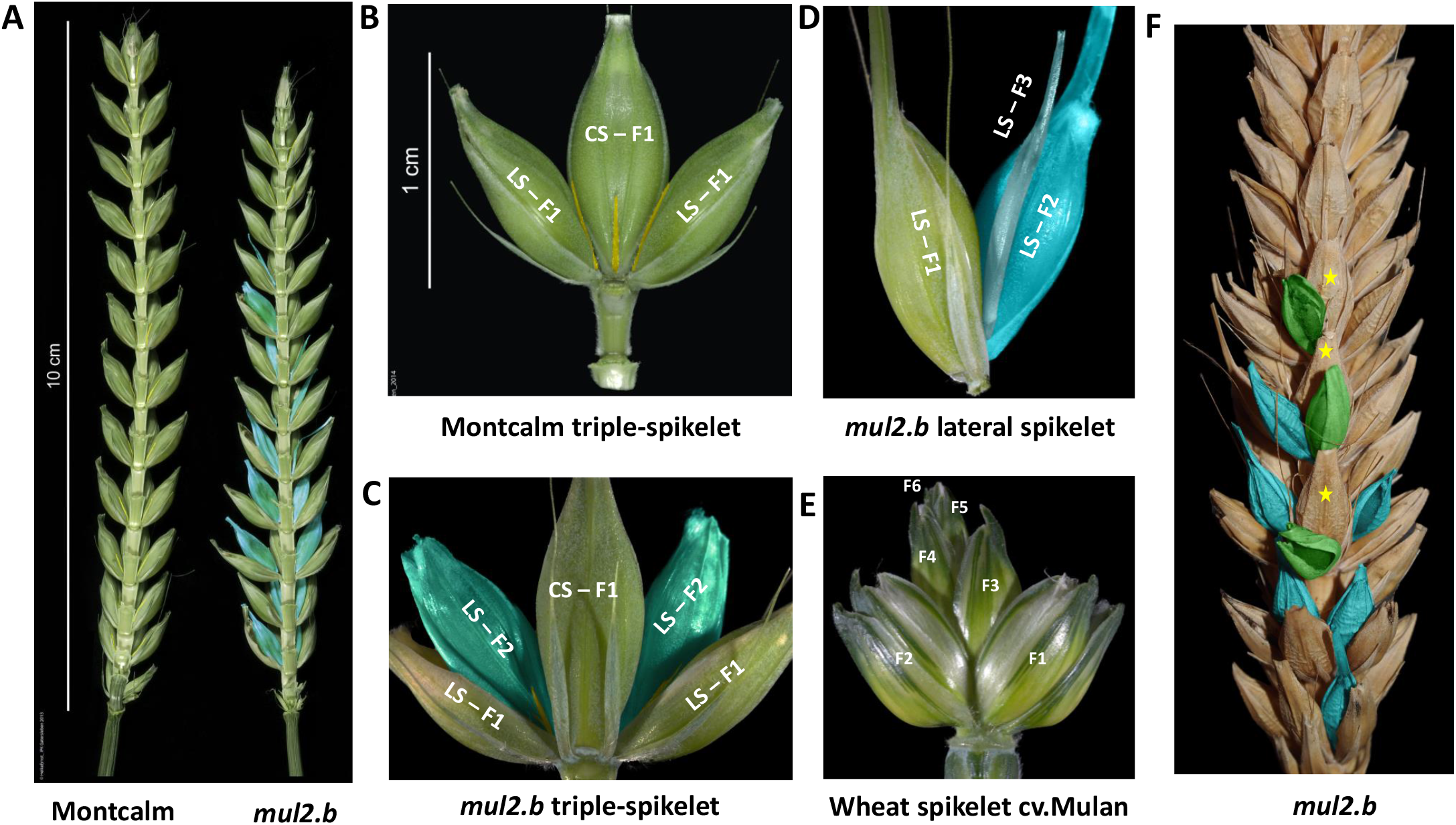
Spike and spikelet phenotypes of cv. Montcalm and derived mutant *mul2.b*. (A) cv. Montcalm and *mul2.b* spikes. The dorsal triple spikelet nodes of Montcalm and *mul2.b* are stripped-off in-order to visualize the rachilla of the lateral spikelets from ventral nodes. The rachilla of lateral spikelets in Montcalm and the second floret on the elongated rachilla of *mul2.b* are colored in yellow and blue respectively. (B) Triple-spikelet node of Montcalm with rachilla of central and lateral spikelets colored in yellow. (C) Triple spikelet node of *mul2.b* mutant showing the formation of the second floret in lateral spikelet nodes (LS-F2). The rachilla on LS-F2 is colored in yellow. (D) The lateral spikelet node of *mul2.b* showing the formation of the second (LS-F2) and third florets (LS-F3). (E) A representative wheat spikelet with six visible florets (F1-F6). (F) *mul2.b* spike showing additional florets in lateral spikelets (blue colored) and supernumerary spikelets (green colored) on the abaxial side of primary central spikelets (indicated by yellow stars). Awns in all images (except 1E) are clipped-off for better visualization of spikelet features.

The genetic control of determinate spikelet fate was studied in rice (*Oryza sativa* L.), wheat, and maize (*Zea mays* L.). In these three species, orthologous APETALA 2 (AP2) transcription factors, such as maize INDETERMINATE SPIKELET1 (IDS1)/TASSELSEED 6 (Chuck et al. 2008; Chuck et al. 2007; Chuck et al. 1998), rice IDS1 (Lee and An 2012), and wheat Q (*AP2L5*) (Debernardi et al. 2017; Greenwood et al. 2017; Simons et al. 2006), were shown to negatively control floret number per spikelet by inhibiting rachilla elongation. The paralog of maize *IDS1*, sister of *IDS1* (*SIDS1*), and rice *SUPERNUMERARY BRACT* also regulate floret number per spikelet by inhibiting rachilla elongation (Chuck et al. 2008; Lee and An 2012; Lee et al. 2007). Interestingly, the corresponding barley ortholog of *IDS1*, i.e., *HvAP2L-H5*, was shown to promote the indeterminate nature of the inflorescence apex and also spikelet determinacy probably by regulating rachilla elongation (Zhong et al. 2021). Apart from the *Q* gene in wheat, the *GRAIN NUMBER INCREASE 1* (*GNI1*) is shown to inhibit rachilla development and growth and thereby regulates floret number per spikelet (Sakuma et al. 2019; Sakuma and Schnurbusch 2020). The *sham ramification 1* (*shr1*) and *shr2* mutants in tetraploid wheat show an elongated rachilla. The genes underlying these loci are not known yet (Amagai et al. 2014).

In rice, genes that potentially regulate spikelet determinacy include *LEAFY HULL STERILE 1* (*OsMADS1* (Jeon et al. 2000)), *MOSAIC FLORAL ORGANS* 1 (*AGAMOUS LIKE 6* (Ohmori et al. 2009)), and *MULTIFLORETSPIKELET1 (AP2/ERF*; (Ren et al. 2013)). Mutations in these genes promote rachilla elongation and extra floret formation rendering spikelets indeterminate. Apart from these, rice genes such as *DOUBLE FLORET1/EXTRA GLUME 1, FLORAL ORGAN NUMBER 4, TONGARI-BOUSHI 1*, and *LATERAL FLORET 1* are also shown to regulate the floret number/spikelet probably through suppression of rachilla elongation (Ren et al. 2019; Ren et al. 2018; Tanaka et al. 2012; Zhang et al. 2017). The results from these studies provided us with a lead that the spikelet determinacy is a plastic trait modulated by the genetic makeup of respective grass species and that the extent of rachilla elongation or suppression potentially defines the floret number per spikelet.

Within Triticeae, barley shows a distinctive spike architecture, imparted by the distichously arranged triple-spikelet units along the rachis. Each spikelet triplet is constituted by one central (CS) and two lateral spikelets (LS), the two- and six-rowed barley types are defined based on the fertility status at the LSs. In two-rowed barley, the CSs are fertile and produce grains, while LSs remain sterile. Whereas in six-rowed, both CSs and LSs are fertile and bear grains. The fertility status of lateral spikelets is predominantly regulated by the *SIX-ROWED SPIKE 1* (*VRS1*, class I HD-ZIP; (Komatsuda et al. 2007)), along with other regulators/modifier genes (Bull et al. 2017; Koppolu et al. 2013; Ramsay et al. 2011; van Esse et al. 2017; Youssef et al. 2017). Thus, the archetypical barley spike possesses one CS positioned on the spike axis with an acute angle, flanked by two LSs positioned slightly away from the spike axis. In wheat, a single spikelet per rachis node is generated with an acute angle to the rachis similar to barley CS. In both wheat and barley, spike mutants that deviate from such a canonical arrangement of spikelets exist that have the potential to generate more spikelets and grains. In barley, a class of mutants called *extrafloret* (*flo*; *flo-a, -b, and -c*) develop either a complete extra or supernumerary spikelet (the locus name *extrafloret* is a misnomer) or rudimentary floral bracts at the base of typical CS on the abaxial side of the same rachis node (Druka et al. 2011; Lundqvist and Franckowiak 2015). The causal genes underlying these loci are not known. A similar supernumerary spikelet (SS) phenotype is also evident in mutants of barley *COMPOSITUM1* (Poursarebani et al. 2020), *COMPOSITUM2* (Poursarebani et al. 2015)) and *VRS4* (Koppolu et al. 2013). The wheat ortholog of barley *COM2* was also shown to regulate SS formation in hexaploid (Dobrovolskaya et al. 2015) and tetraploid branched miracle wheat (*TtBH^t^*; (Sakuma and Schnurbusch 2020; Wolde et al. 2019). Apart from these genes, the SS phenotype in wheat is also conditioned by the photoperiod-dependent floral induction gene, *PHOTOPERIOD 1* (*PPD1*), and the axillary meristem growth regulator *TEOSINTE BRANCHED 1* (*TB1*). Both *PPD1* and *TB1* regulate SS formation by regulating *FLOWERING LOCUS T1* (*FT1*) that induces a cascade of floral meristem identity genes important for floral induction (Boden et al. 2015; Dixon et al. 2018).

In this study, we investigated the inheritance of spikelet determinacy (floret number per spikelet) and supernumerary spikelet phenotypes in a barley *X*-ray induced mutant *“multiflorus2.b”* (*mul2.b;* progenitor–Montcalm). The barley *mul2.b* mutants produce non-canonical multi-floreted, indeterminate spikelets with elongated rachillae, a feature reminiscent of wheat spikelets. Moreover, our in-depth mutant phenotyping and allelism analyses show that *mul2.b* also induces the supernumerary spikelet phenotype. Based on our analyses using exome capture-based bulked-segregant analysis, genetic and QTL mapping studies, we reveal that the multi-floreted and supernumerary spikelet phenotypes show quantitative inheritance patterns.

## Materials and Methods

### Plant material and growth conditions

All experiments reported in this study were conducted at the Leibniz Institute of Plant Genetics and Crop Plant Research, Gatersleben, Germany. Barley genotypes cv. Montcalm, cv. Morex, *multiflorus2.b* (*mul2.b*; GSHO 1394) – a betatron induced *X*-ray mutant of Montcalm, and Bowman Near Isogenic line mutant *extra floret.a* – BM-NIL(*flo.a*), were used for various analyses conducted in this study. The majority of experiments were conducted under greenhouse conditions at the IPK in the period between 2014 and 2020. For greenhouse experiments, initially, barley grains were germinated in 96 well planting trays and grown at 16h/8h (light/dark; photoperiod), and, 20°C/16°C (light/dark; temperature) for two weeks. After collecting leaf material for DNA extraction, the plants were vernalized at 4°C for four weeks and then acclimatized at 15 °C in the greenhouse for a week. Later the plants were potted into 14 cm (Ø) pots and grown at 16h/8h (light/dark; photoperiod), and, 20° C/16° C (light/dark; temperature) until maturity.

Standard practices for irrigation, fertilization, and control of pests and diseases were followed. The genotypes Montcalm and *mul2.b* were evaluated under field conditions in 2015 and 2016 at IPK for the grain morphometry parameters. Each genotype was grown in three independent plots measuring 1 × 1.5 m^2^, with a sowing density of 300 grains/m^2^. Each plot consisted of six rows with 20 cm spacing between rows.

### Mapping populations and phenotyping

For genetic mapping of the traits lateral spikelet indeterminacy (LS-IN) and central spikelet supernumerary spikelet (CS-SS) phenotypes, the mutant parent *mul2.b* was crossed with Morex and Montcalm to generate two independent mapping populations, Morex × *mul2.b* and *mul2.b* × Montcalm. The F_1_ hybrids were selfed and the resulting F_2_ individuals and F_2:3_ families were used for phenotypic and genotypic analyses. To test the allelism between *mul2.b* and BM-NIL(*flo.a*), reciprocal crosses were made between the two mutants, and the CS-SS phenotype were analyzed at F_1_ and F_2_ generations.

The Morex × *mul2.b* (POP 2014-1; 163 F_2s_) and *mul2.b* × Montcalm (POP 2014-2; 130 F_2s_) populations were initially phenotyped in 2014 under greenhouse conditions. The LS-IN phenotype was scored as strong or moderate to weak and wildtype based on visual observation of rachilla elongation or extra floret phenotype in all spikes of each F_2_ plant after grain filling. F_2_ plants with strong phenotype showed approximately 75% of spikes with a second floret that either produced a grain or remained empty. Whereas the F_2_ lines that showed elongated rachilla or improperly formed second floret were scored into weak to moderate phenotype class. The CS-SS phenotype in central spikelets was quantified by counting the number of occurrences of extra spikelets or rudimentary bract-like structures on all spikes of each F_2_ plant.

In 2015, we phenotyped Morex × *mul2.b* population again under greenhouse conditions (136 F_2s_; POP 2015) by quantifying both LS-IN and CS-SS phenotypes. For quantification, of LS-IN phenotype we classified the phenotype into *mul2.b class-I, -II, -III*, and -*IV* based on the additional floret or rachilla elongation phenotype (please see results section for the description of each phenotypic class). The number of instances of *mul2.b-I, -II, -III*, and -*IV* phenotypes were recorded for all lateral spikelet positions on spike-bearing tillers of each F_2_ plant. Similarly, the number of instances of supernumerary spikelets born from the central spikelet region was counted.

In 2016, a relatively larger F_2_ population of Morex × *mul2.b* was further phenotyped under greenhouse conditions (179 F_2s_; POP 2016) for the LS-IN and CS-SS phenotypes. The phenotype quantification data from POP 2016 was used for whole-genome QTL mapping.

For scoring the CS-SS trait in F_2s_ of BM-NIL(*flo.a*) × *mul2.b*, the phenotype was classified into six classes. These include WT (no phenotype), very weak (one spike with CS-SS phenotype out of all spikes/plant), weak (~10-15% of spikes with CS-SS phenotype), moderate (~25% of spikes with CS-SS phenotype), strong (~50% of spikes with CS-SS phenotype), and very strong (~75-100% of spikes with CS-SS phenotype.)

### Grain morphometry analysis of Montcalm and mul2.b

To evaluate grain morphometry parameters of Montcalm and *mul2.b*, spikes from the field (2015 and 2016) and greenhouse (2020) grown plants were hand-harvested and threshed. Representative grain samples from each genotype (3–4 grain samples/plot/genotype) were analyzed for the thousand-grain weight (TGW), grain area (mm^2^), grain width (mm), and grain length (mm) using a digital seed analyzer/counter Marvin (GTA Sensorik GmbH, Neubrandenburg, Germany). From the greenhouse experiment (2020), the grain morphometry parameters were collected for central and lateral spikelet grains independently. For this analysis, central and lateral spikelet grains of twelve independent plants from each genotype were used. Apart from these, traits such as plant height, total productive and unproductive tillers were scored from the greenhouse experiment (2020).

### Phenotyping of Montcalm and mul2.b immature spikes

Immature spikes of Montcalm and *mul2.b* genotypes were phenotyped from Waddington stage W2.0 (double ridge) to W6.0 (Waddington et al. 1983) to follow rachilla development in lateral spikelets. The spike meristems from respective genotypes were hand dissected and staged under a stereomicroscope (Zeiss Stemi 2000-C; Carl Zeiss™). The maximum yield potential in Montcalm and *mul2.b* (number of rachis nodes per spike * three spikelets per node) was counted on the main culm spike approximately at W8.0.

### Scanning electron microscopy

Scanning electron microscopy (SEM) was performed on immature spikes that are approximately at W4.0, W4.5, W5.0, W5.5, and W6.0 (central spikelet staging) from greenhouse-grown plants. SEM was conducted following an established protocol at IPK (Lolas et al. 2010; Poursarebani et al. 2020).

### Exome sequencing of Morex × mul2.b – POP 2014-1

For exome sequencing, the DNAs from 25 wild type and 27 mutant F_2_ lines were bulked from Morex × *mul2.b* population grown in 2014. The two DNA bulks were subjected to exome capture and sequencing according to Mascher et al. (2013). Read mapping, allele frequency visualization, and read depth analysis of exome sequencing data of mutant and wild-type bulks were performed as described in Mascher et al. (2014) using the POPSEQ map.

### Marker development, linkage, and QTL analysis on 2H and 6H chromosomes

The LS-IN and CS-SS phenotype data from Morex × *mul2.b*–2015 (POP 2015) was used for marker and phenotype linkage analysis on 2H and 6H barley chromosomes. For mapping, the polymorphic SNPs were selected based on exome capture SNPs from chromosomes 2H and 6H where allele frequency differences between mutant and wild type bulks reached their maximum. On chromosome 2H, 11 SNPs were selected in the interval 33.70–87.70 cM (Morex POPSEQ positions). On 6H, seven SNPs were selected in the interval 31.90–92.20 cM. Selected SNPs were converted to restriction enzyme-based Cleaved Amplified Polymorphic Sequence (CAPS) markers (**Supplementary Table 1**) using Neb cutter v2.0 (New England Biolabs Inc). Restriction digestion was performed according to the manufacturer’s protocols. Resulting DNA fragments were separated on 2% agarose gels for genotyping. Genotypic and phenotypic data were subjected to linkage and QTL analyses in Joinmap 4.1 and Genstat 19 respectively.

### GBS library construction, sequencing, read alignment, and variant calling

Total genomic DNA from the F_2_ lines of Morex × *mul2.b* (POP 2016) was extracted as described in Poursarebani et al. (2020). Genomic DNA was digested with *PstI* and *MspI* restriction enzymes and processed for GBS library construction as described in Wendler et al. (2014). Individually barcoded samples were pooled in equimolar concentrations and sequenced on the Illumina HiSeq 2500 (single-read, 107 cycles), using custom sequencing primer (Wendler et al. 2014) as given in manufacturers protocol (Illumina Inc).

GBS reads were trimmed using cutadapt (Martin 2011) and aligned to the reference genome sequence of Morex (Mascher et al. 2017) with BWA-MEM v0.7.12a (Li 2013). Alignment files were converted to binary alignment map (BAM) format with SAMtools (Li et al. 2009), sorted by reference position, and indexed with NovoSort (Novocraft Technologies Sdn Bhd, Malaysia, http://www.novocraft.com/). Variants were called using SAMtools/BCFtools v1.3 specifying the option ‘-DV’ while running SAMtools mpileup (Li 2011)), to obtain the per-sample number of non-reference reads. An AWK script (available at https://bitbucket.org/ipk_dg_public/vcf_filtering) was used to filter the Variant Call Format (VCF) output file and call genotypes based on the ratio between the depth of the alternative allele and the total read depth (DV /DP). A minimum mapping quality score of 40 was applied. The SNP matrix was created applying the following threshold: a minimum read depth of 2X per genotype and an overall maximum fraction of missing calls of 80%. Only biallelic SNPs were considered. Data for SNPs was filtered with ≤10% missing data points, two or more linked markers showing identical marker scores across all F_2_ individuals (no recombination), and variants showing distorted marker scores. The final variant set was composed of 605 markers and was used for genetic linkage mapping.

### Whole-genome linkage mapping with GBS markers

Genetic linkage maps were constructed with data from 605 GBS markers. Markers were initially grouped according to the SNP physical position assignment based on Morex reference. Markers from each group were then ordered using the maximum likelihood mapping algorithm and default linkage parameters in Joinmap 4.1. Recombination frequencies between markers were converted to map distances by the Kosambi mapping function.

### Phenotypic data analyses of Morex × mul2.b–POP 2016

Phenotypic data for LS-IN and CS-SS traits were collected from up to 12 tillers. To calculate the individual variance components of the genotype, tillers, and residuals, the following linear mixed-effect model was used by assuming all effects except the intercept as random:

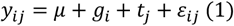

where, *y_ij_* is the phenotypic value of the *i^th^* genotype in the *j^th^* tiller, *μ* is the common intercept term, *g_i_* is the effect of the *i^th^* genotype, *t_j_* is the effect of the *j^th^* tiller, and *ε_ij_* is the corresponding residual term as 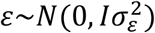 with *I* and 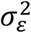 being the identity matrix and residual variance, respectively. The repeatability (*H*^2^) among the tillers was accordingly calculated as:

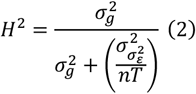

where, σ_g^2 and σ_ε^2 denote the variance components of the genotype and residuals, respectively; nT represent the number of tillers. The best linear unbiased estimations (BLUEs) across tillers for a given genotype were calculated by assuming the intercept and genotype effects as fixed in eq. 1. It should be noted that we evaluated the traits in an F_2_ population; therefore, by necessity, we collected the phenotypic data from single plants. The calculations mentioned above were performed to (i) observe if there exists a large and significant (P<0.001) variance for the spike bearing tillers, and (ii) normalize the phenotypic value of the investigated traits per spike since it gives a more meaningful view of the studied traits. The BLUEs were plotted in the form of histograms. To observe the genetic nature of the investigated traits, the Shapiro-Wilk test was performed to monitor if the data were normally distributed at P<0.05. Moreover, to observe the genetic relationship among the investigated traits, Pearson’s product-moment correlation was calculated based on the BLUEs as:

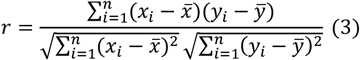

where *x* and *y* denote the BLUEs of the studied traits. Unless stated otherwise, all calculations were performed in software R (R core team) using package lme4 (Bates et al. 2015).

### Whole-genome QTL mapping in Morex × mul2.b–POP 2016

The BLUEs from LS-IN (*mul2.b-I, -II, III*, and -*IV*) and CS-SS phenotypes and 605 GBS marker data were used for QTL mapping in Genstat 19. Initially, simple interval mapping (SIM) was conducted with single environment, single trait linkage analysis. A step size of 5 cM, genome-wide significance level (alpha) of 0.05, and minimum co-factor proximity of 50 cM were used for QTL detection. The minimum separation distance for two consecutive QTLs was set at 30 cM. The co-factors identified from SIM were used for QTL analysis by composite interval mapping (CIM). This process was repeated until no new QTLs were detected. The final QTL model was selected based on background selection (significance 0.05) on the selected co-factors. QTL boundaries (lower and upper), additive, dominance effects, and phenotypic variation explained by the QTL were defined by using the final QTL model. The procedure was repeated for QTL detection in all traits scored. Genetic linkage maps and QTL data were graphically represented using MapChart 2.2 (Voorrips 2002).

### Phenotypic analysis of LS-IN and CS-SS in F_2:3_

To validate the genetic and phenotypic effects among QTL loci identified in F_2_, we performed the phenotypic analysis in F_2:3_ for LS-IN and CS-SS phenotypes. For this purpose, we have generated an F_2:3_ population comprising 1180 individuals segregating for Morex and *mul2.b* alleles at 2H and 6H QTLs. The F_2:3_ lines were genotyped with at least four CAPS markers linked to the 2H (M269389, M5729, 2560790, and M49460) and two markers linked to 6H (FL277086, and FL42060) QTLs. All F_2:3_ lines were phenotyped for floret number in LSs and SS phenotype in CSs. The combinations of 2H and 6H allele frequencies and the corresponding phenotypic values for the respective genotypes are plotted as box plots. All box plots in this study were generated using BoxPlotR (Spitzer et al. 2014).

## Results

### Lateral spikelet determinacy is relaxed in the multiflorus2.b

Spikelets of barley are commonly determinate structures with a single floret being produced per spikelet on its axis i.e. rachilla. After the first floret formation, the elongation of the rachilla is stopped and the rest of the rachilla remains as a thin, pale yellow colored suppressed structure adaxial to the floret (positioned between rachis and palea of the floret; **Fig. 1A, B**). This scenario applies to both CS and LSs of all spikelet triplets along the inflorescence axis. However, in the LSs of *mul2.b* mutants, upon initiation of the first floret (LS-F1), the rachilla further elongates which leads to the development of a second floret (LS-F2) at the articulation junction of the elongated rachilla (**Fig. 1A, C**). In *mul2.b* mutants, the elongated rachilla becomes visible on the adaxial side of the second floret (LS-F2; **Fig. 1C**). Often the elongated rachilla on the second floret of *mul2.b* LSs is enlarged, pale green-colored, indicating that the rachilla elongation is not completely abolished after the second floret formation. In line with this, the rachilla in *mul2.b* LS occasionally elongates beyond the second floret and bears a third floret (LS-F3; **Fig. 1D**). Thus, the rachilla elongation suppression in LSs appears to be lost in *mul2.b* rendering the lateral spikelets indeterminate, a feature reminiscent of indeterminate wheat spikelets (**Fig. 1E**). The rachilla elongation and formation of second and third florets were predominantly restricted to the basal half of the spike (**Fig. 1A**)

To understand the developmental events during the rachilla primordium initiation and elongation, we undertook scanning electronic microscopy (SEM) analysis initially in wild-type progenitor Montcalm. The general barley spike meristem developmental progression starts with the transition of vegetative shoot apical meristem into reproductive inflorescence meristem (IM). The IM initiates spikelet ridge meristem that further differentiates into one central (CSM) and two lateral spikelet meristems (LSM), followed by the formation of two glume primordia in the flanks of each spikelet meristem (Koppolu and Schnurbusch 2019). At this point, in Montcalm we observed that the floret meristem (FM) first differentiates lemma primordium on the abaxial side. After the initiation of lemma primordium, the adaxial side of the floral meristem buds-off to initiate the rachilla primordium (**Fig. 2A**). Later the residual floral meristem is consumed into the initiation of lodicule, stamen, and carpel primordia (**Fig. 2B-E**). The turn of meristem differentiation events is identical in both CSM and LSM. However, visualizing rachilla primordium in CSM is technically challenging especially at the very early stages of spike meristem development due to the acute angle between CSM and the axis of the spike meristem. Hence, we followed the rachilla primordium development in LSs that are positioned away from the spike meristem axis at appropriate floral primordia initiation time points. In general, the organ primordia differentiated from the floret meristem of Montcalm tended to grow normally with an apparent increase in size over the immature spike developmental phases analyzed (**Fig. 2A-E**). However, the rate of rachilla primordium growth appeared to be slowed down compared to other organ primordia differentiated from the FM (**Fig. 2A-E**). At around W5.0 (central spikelet staging), the rachilla is barely visible in the lateral spikelets at the base and middle of the spike (the developmentally advanced portion of the spike; **Fig. 2C**). Later on, during the spike growth period, the rachilla ceases to grow and remains as a suppressed structure in Montcalm (as seen in **Fig. 1A, B**). In the *mul2.b* mutant, the rachilla primordium was initiated after the differentiation of lemma primordium similar to Montcalm (**Fig. 2F**). Nevertheless, the rachilla primordium of *mul2.b* LSs was not retarded in development and continued to increase in size along with other floral organs differentiated from floret meristem (**Fig. 2F-K**). Over different phases of immature spike development, the rachilla primordium elongated further to form a second FM (**Fig. 2I**). Importantly, the rachilla elongation and formation of additional FM in *mul2.b* was restricted to the LSs; whereas, the rachilla of CS remained as a suppressed structure (**Fig. 2K**).

**Figure 2.**
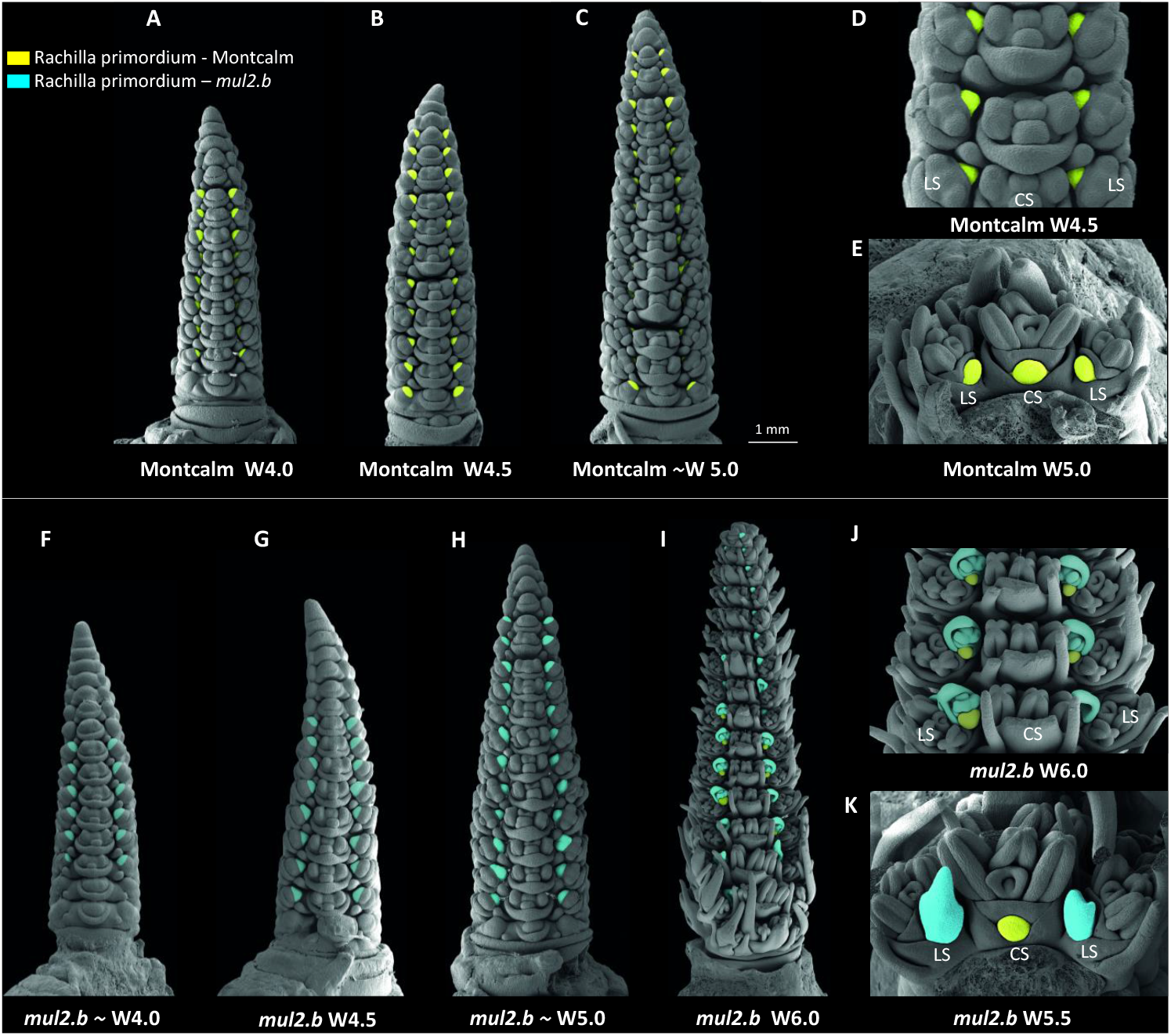
Scanning electron micrographs highlighted with rachilla primordia in cv. Montcalm and derived mutant *mul2.b*. (A-C) Immature spikes of cv. Montcalm at Waddington W4.0 (A), W4.5 (B), and ***~***W5.0 (C). (D) Close-up view of three triple-spikelet nodes of Montcalm (E) Top-down view of single triple-spikelet of Montcalm. The lateral spikelet rachilla primordia in figures 2A-E are colored in yellow. (F-I) Immature spikes of *mul2.b* at ***~***W4.0 (F), W4.5(G), ***~***W5.0 (H), and W6.0 (I). (J) Close up view of three triple-spikelet nodes of *mul2.b*. (K) Top-down view of single triple-spikelet of *mul2.b*. The rachilla primordium/second floret in lateral spikelets in figures 2F-K are colored in blue. The rachilla primordium seen on the second floret in 2I, 2J, and central spikelet in 2K is colored in yellow. All immature spikes in figure 2 are staged based on the central spikelet development.

Since the LSs of *mul2.b* mutants produced additional florets that are frequently fertile and grain-bearing, we evaluated various yield and biomass-related traits in *mul2.b* and Montcalm. Interestingly, the immature spikes of *mul2.b* showed a significantly increased maximum yield potential compared to Montcalm (Median spikelet number at W8.0–Montcalm 88.5 spikelet primordia; *mul2.b* 102 spikelet primordia; p-value: 5.07E-13). However, the number of fertile spikelets was significantly lowered in *mul2.b* (Median fertile spikelets – 39 in *mul2.b*, 45 in Montcalm; p-value: 0.001) probably owing to the enhanced sink strength in the basal half of the spike and consequent post-anthesis growth abortion of spikelets in the apical part of the spike (**Fig. S1A-C**). The grain parameters such as thousand-grain weight (TGW), grain area (GrA), width (GW) and length (GL) did not show significant differences between Montcalm and *mul2.b* when grains from CS and LS were analyzed together (**Fig. S2A-D**). However, the differences for these traits became apparent between Montcalm and *mul2.b* when the CS and LS grains were measured separately. Interestingly the TGW of CS grains was significantly increased in *mul2.b* (p-value: 0.020) due to the slight increase in grain area, mainly contributed by the increase in GW (**Fig. S2E-H**). The TGW of LSs did not show a significant difference between Montcalm and *mul2.b* (**Fig. S2I**). The parameters GW and GL followed opposite tendencies in *mul2.b* LSs where GW was slightly higher and GL was significantly lowered (p-value: 3.61E-05) compared to Montcalm, probably leading to similar GrA and TGW (**Fig. S2I-L**). Apart from the grain parameters, the plant height was significantly reduced in *mul2.b* (p-value: 0.020), whereas the number of tillers including productive spike-bearing tillers and unproductive tillers remained the same (**fig. S1D-G**).

### Lateral spikelets of mul2.b display a gradual rachilla elongation phenotype

The extent of rachilla elongation and the type of additional structures being formed in the LSs of *mul2.b* varied among spikelets of each spike. To quantify these additional structures, we devised a score to classify them into four phenotype component classes such as *mul2* – *class I* (*mul2-I*), *mul2-II, mul2-III*, and *mul2-IV*. The *mul2-I* had the strongest LS rachilla elongation phenotype with the formation of a second floret that is often fertile and an occasional third floret (**Fig. 3B, C,** and **Fig. S3B**). Also, the second floret with an elongated awn is part of the *mul2-I* class. The reproductive organ development is mostly complete in *mul2-I* except for the occasional lack of carpel growth. The second floret of *mul2-II* is comparatively smaller without awn elongation (**Fig. 3D, E** and **Fig. S3C**). Occasionally, the second floret in the *mul2-II* class had stamens but carpel growth is completely abolished. In the *mul2-III* class, the unsuppressed rachilla showed bi- or trifurcation at the base that further elongated into filament-like structure (**Fig. 3F, G** and **Fig. S3D**). The *mul2-IV* phenotypic class is similar to *mul2-III* except that the unsuppressed rachilla did not elongate into a filament-like structure (**Fig. 3H, I** and **Fig. S3E**). In both *mul2-III*, and -*IV* the elongated rachilla did not attain a second floret identity as seen in *mul2-I* and *mul2–II*. The spatial occurrence of *mul2-I, -II, -III, -IV* individual component classes did not show any preferential pattern in the spike indicating the random nature of their occurrence along the spike axis (**Fig. S4**). Throughout this report, we referred to the rachilla elongation phenotype of LSs as lateral spikelet indeterminacy (LS-IN).

**Figure 3.**
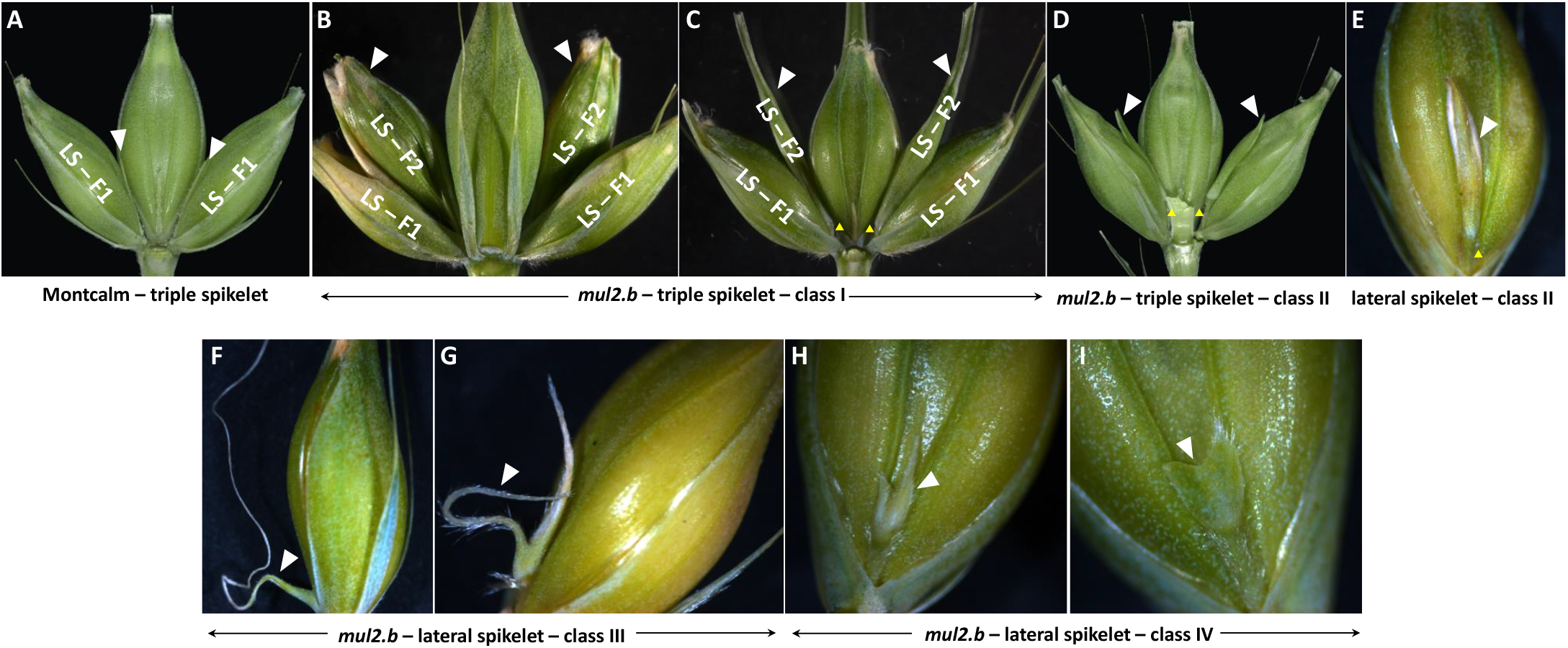
Phenotype classification for lateral spikelet indeterminacy. (A) Representative spikelet-triplet showing wild-type rachilla of lateral and central spikelets. (B-E) elongated lateral spikelet rachilla developing class I (B, C) and class II additional florets (D, E). (F-G) Class III rachilla elongating into a thin and long filament-like structure. (H-I) Class IV phenotype showing a bifurcated (H) or wedge-shaped rachilla in lateral spikelets (I). Rachilla/additional florets/rachilla malformations in lateral spikelets are indicated with white arrowheads. Yellow arrowheads in figure 3D and E indicate a pedicel below the extra floret in lateral spikelets.

### Supernumerary spikelet formation in central spikelets of mul2.b

In contrast to the loss of spikelet determinacy in the LSs, the *mul2.b* mutant showed another interesting feature: the formation of supernumerary spikelets (SS) exclusively in the CS. The SSs were formed abaxial to the typical CSs (**Fig. 1F**). The observed supernumerary spikelet phenotype in *mul2.b* ranged from the formation of rudimentary floral bract-like structures to complete spikelets with occasional grain formation (**Fig. 1F**). Such a SS phenotype is also evident in the barley mutant *extrafloret.a* (BM-NIL(*flo.a*). Interestingly, the SSs that originated at the CSs were occasionally indeterminate with rachilla elongation and formation of second floret similar to the indeterminate LS phenotype in *mul2.b* (**Fig. 1F**). In this report, we termed the SS phenotype on the CS as “CS-SS”.

### Genetic mapping identifies genomic regions responsible for the LS-IN and CS-SS phenotypes

Previous studies reported that the LS-IN trait in *mul2.b* showed a monogenic recessive inheritance pattern (Walker et al. 1963). Hence, we developed two F_2_ mapping populations by crossing *mul2.b* with Morex (Morex × *mul2.b*; POP 2014-1) and Montcalm (*mul2.b* × Montcalm; POP 2014-2) for genetic mapping. Of the 163 F_2_ lines phenotyped from POP 2014-1, 35 and 83 F_2s_ showed strong and weak to moderate LS-IN phenotypes, respectively, while 45 F_2s_ were WT, suggesting that the observed phenotype segregation was in contradiction with the previously proposed monogenic recessive inheritance pattern for the LS-IN trait (Walker et al. 1963). From our phenotypic analysis, the LS-IN phenotype appeared to follow an incomplete dominance with homozygous mutant F_2s_ showing strong and the heterozygous F_2s_ showing weak to moderate LS-IN phenotypes (χ^2^ = 0.591, *p-value* = 0.442). A similar incomplete dominant inheritance was also observed for the POP 2014-2 (**Supplementary Table 2**). With respect to the CS-SS phenotype, 99 F_2s_ of POP 2014-1 showed the phenotype whereas 64 F_2s_ did not show a phenotype; and hence, were wild type for CS-SS. The majority of the F_2s_ with CS-SS phenotype also had the LS-IN phenotype (70 out of 99 F_2s_), while 29 F_2s_ that were wild type for LS-IN also showed a weak CS-SS phenotype. Thus, the phenotypic inheritance analysis in F_2s_ indicated that the *mul2.b* phenotype is in fact a combination of LS-IN and CS-SS phenotypes.

For identifying the genomic region(s) harboring the *mul2.b* mutation, we adopted sequencing-based bulked-segregant analysis (BSA) by exome capture (Mascher et al. 2014; Mascher et al. 2013). To this end, DNAs from 27 mutant F_2s_ with a strong LS-IN phenotype, of which 20 individuals also had the CS-SS phenotype (**Figure 4A**), and 25 wild-type F_2s_ were pooled to form the LS-IN/CS-SS mutant and wild-type bulks. The two bulks were subjected to exome capture and subsequently sequenced on Illumina Hiseq2000. The single nucleotide polymorphisms were identified by mapping sequence reads onto the Morex reference genome. The SNP allele frequencies from the two pools were visualized along the barley physical and genetic maps. Surprisingly, the allele frequency distribution showed two distinct peaks on chromosomes 2H and 6H where in both cases the mutant allele frequencies for the combined LS-IN/CS-SS phenotype increased over 75% and wt allele frequency dropped to about 25% (**Fig. 4B**).

**Figure 4.**
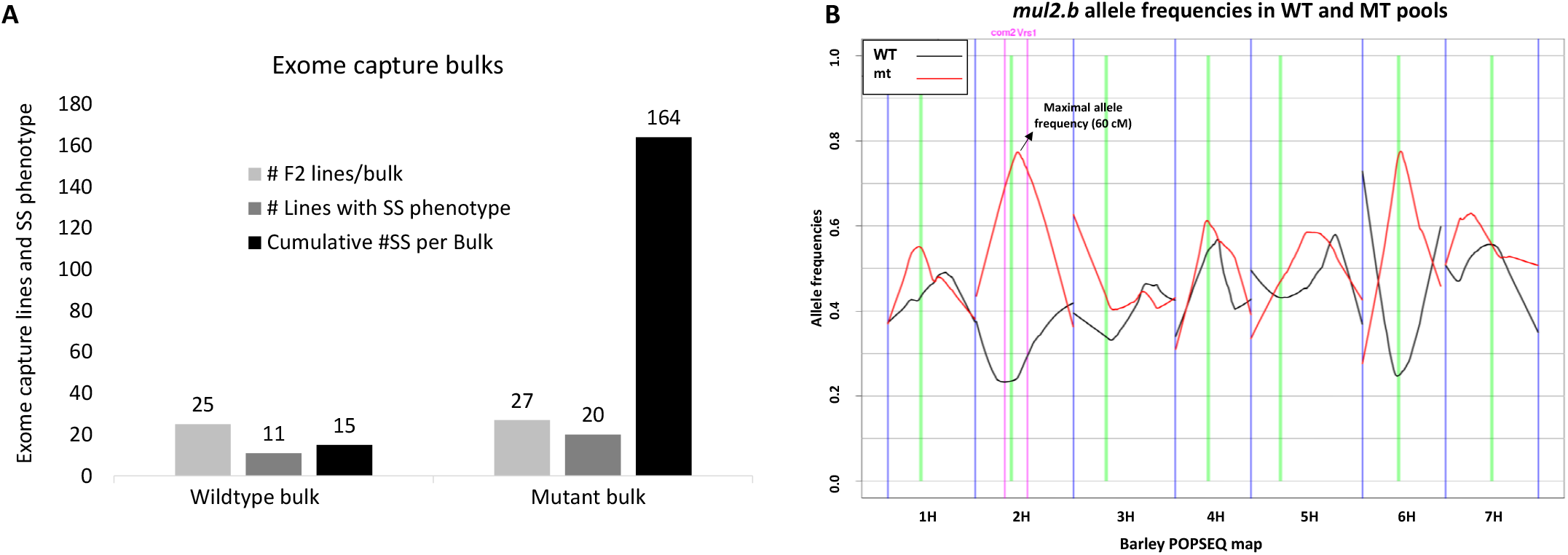
Exome capture-based bulked segregant analysis in Morex × *mul2.b* (POP 2014-1). (A) Graph showing number of F_2_ individuals per exome capture pool, number of exome capture F_2_ individuals showing supernumerary spikelet phenotype, and strength of supernumerary spikelet phenotype (total supernumerary spikelets per pool) in WT and mt pools. (B) Allele frequency plot showing the frequency of wild-type (Morex) and mutant (*mul2.b*) allele distributions across seven barley chromosomes.

### A novel locus regulates the CS-SS phenotype in mul2.b

In our BSA, allele frequencies peaked on 2H and 6H, indicating that the combined phenotypes of LS-IN and CS-SS are regulated by two independent loci in *mul2.b*. Previous genetic analyses linked the *flo.a* phenotype on the short arm of 6H (Druka et al. 2011). Interestingly, the CSs of the barley BM-NIL(*flo.a*) mutant showed the identical CS-SS spikelet phenotype (Lundqvist and Franckowiak 2015) as *mul2.b* (**Fig. 5A-B**), providing the possibility that both loci might be identical. To examine this idea (**Fig. 5A-B**), we made reciprocal crosses between these two mutant loci to evaluate allelism between them. The CS-SS phenotype in *flo.a* was previously shown to follow a monogenic recessive inheritance (Gustafson 1969). The F_1s_ generated from the *mul2.b* and BM-NIL(*flo.a*) crosses showed a strong CS-SS phenotype (**Fig. 5A-C**), potentially suggesting that the CS-SS phenotype in both BM-NIL(*flo.a*) and *mul2.b* is regulated by a single locus. However, evidence can only be provided by checking phenotypes of F_2s_. We therefore evaluated the F_2_ generation of the BM-NIL(*flo.a*) × *mul2.b* cross for the CS-SS phenotype to further verify the potential allelic relation of *flo.a* and *mul2.b*. Surprisingly, we found segregation for the CS-SS phenotype among the F_2_ progenies: i.e. of the 395 F_2s_ analyzed, 106 did not show the CS-SS phenotype, whereas 289 F_2s_ showed a very weak to very strong CS-SS phenotype (**Fig. 5D**). The observed phenotype segregation in F_2_ thus indicated that the CS-SS phenotype in *mul2.b* and *flo.a* is most likely regulated by two-independent loci (**Fig. 5E**). The allele and phenotype proportion analysis for the BM-NIL(*flo.a*) × *mul2.b* dihybrid cross showed that the 6H locus regulating the CS-SS phenotype in *mul2.b* followed a recessive inheritance (χ^2^ = 3.859, *p-value* = 0.0495). From our exome capture data, the highest allele frequency difference for the 6H peak lied between 59.92 – 61.05 cM (POPSEQ position), whereas the *flo.a* locus was mapped to 52.20 cM (data not shown), indicating that the CS-SS phenotypes in *mul2.b* and *flo.a* are regulated by two independent loci on 6H.

**Figure 5.**
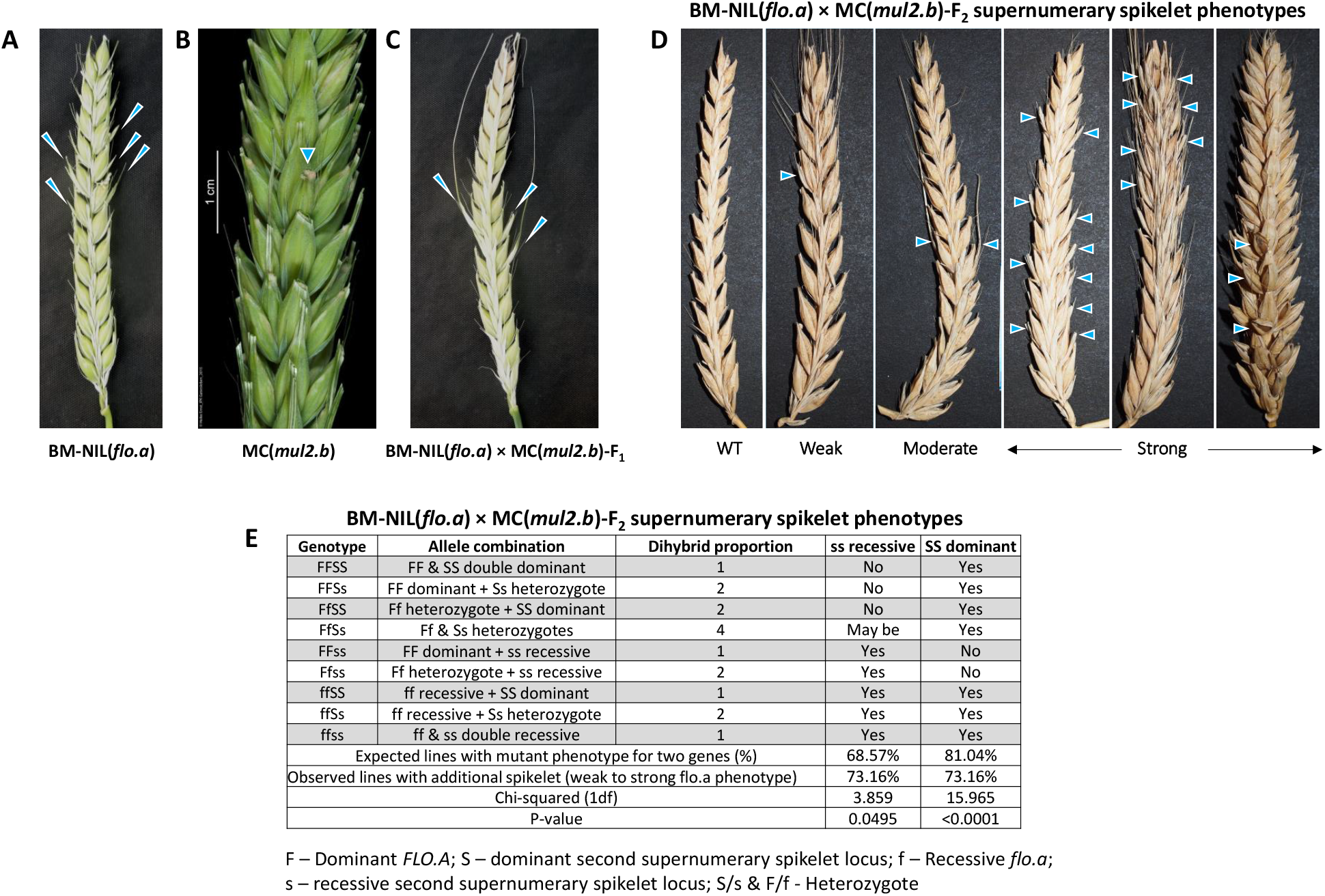
Allelism analysis between *mul2.b* and BM-NIL(*flo.a*). (A-C) Supernumerary spikelet formation in BM-NIL(*flo.a*) (A), *mul2.b* (B), and the respective F_1_ hybrid (C). (D) Supernumerary spikelet phenotypes in BM-NIL(*flo.a*) × *mul2.b* F_2_ individuals. (E) Mode of inheritance of supernumerary spikelet phenotype in BM-NIL(*flo.a*) × *mul2.b*. Supernumerary spikelets are indicated by blue arrowheads. F – Dominant *FLO.A*; S – dominant second supernumerary spikelet locus; f – Recessive *flo.a*; s – recessive second supernumerary spikelet locus; S/s & F/f – Heterozygotes.

### Genetic linkage of the LS-IN and CS-SS phenotypes to 2H and 6H

To conduct a rough genetic linkage analysis of the LS-IN and CS-SS phenotypes we selected 11 SNPs on 2H (33.70 – 87.70 cM POPSEQ) and seven SNPs on 6H (31.90 – 92.20 cM POPSEQ) based on the introgressions identified from BSA. The SNPs were converted to restriction enzyme-based CAPS markers for genotyping. For SNP marker mapping, we phenotyped 136 new F_2s_ of the Morex × *mul2.b* population in the year 2015 (POP 2015). We knew that the barley spike-branching gene *COM2* is also located on 2H (45.0 cM), mutants of which often displayed the LS-IN phenotype (McKim et al. 2018; Poursarebani et al. 2015). To check if the LS-IN phenotype is regulated by *COM2*, we developed a CAPS marker out of *COM2* for mapping in our population. Different from the phenotyping method followed for POP 2014-1 and POP 2014-2, we quantified the LS-IN phenotype in POP 2015, based on *mul2-I, -II, -III*, and -*IV* classes (described earlier in results). In the same F_2s_, we quantified the CS-SS phenotype. Out of 136 F_2s_ phenotyped, 67 showed various quantities of *mul2-I, -II, -III, -IV* LS-IN phenotypes, while 69 were wildtype for LS-IN. This segregation did not fit the incomplete dominant inheritance (χ^2^ = 48.039, *p-value* = <0.0001) observed in POP 2014-1 and POP 2014-2, sugesting some sort of plasticity for the LS-IN phenotype. With respect to the CS-SS phenotype, 62 F_2s_ of POP 2014-1 showed the CS-SS, the majority of which also had LS-IN phenotype while 74 were wt for CS-SS.

For linkage analysis of the LS-IN phenotype, we converted the phenotype quantities (cumulative frequencies of the *mul2-I, -II, -III*, and -*IV* classes) into binary scores along with SNP marker genotypes. However, after comparing the SNP and phenotype marker scores, six out of 136 F_2s_ had a mismatch between phenotype and marker genotypes. These included, two F_2s_ that showed the LS-IN phenotype but had homozygous WT genotype calls and four F_2s_ that were phenotypically wt for LS-IN but had mutant genotype calls at the majority of markers mapped on 2H (**Fig. 6A**). The two F_2s_ that had wt genotype calls on 2H showed a weak LS-IN phenotype (*mul2-III* and -*IV*; **Fig. 6A**). However, performing linkage analysis of the LS-IN phenotype and genotype scores by excluding the six conflicting F_2s_, linked LS-IN in between markers M1865727 and M5729 away from the *COM2* gene (**Fig. 6B**). Since the LS-IN phenotype was measured in a quantitative manner, we were also able to perform QTL analysis for the LS-IN phenotype (cumulative *mul2.b-I, -II, -III, and -IV*) together with CAPS marker data from 2H and 6H. Expectedly based on our previous exome capture data, LS-IN showed a major QTL on 2H with marker M5729 explaining the highest phenotypic value (**Fig. 6C, E**). However, we also noticed another major QTL for the LS-IN phenotype on 6H with FL209450 explaining the highest phenotypic value (**Fig. 6C, E**). Furthermore, our QTL analysis of the CS-SS phenotype with marker data from 2H and 6H showed a single major QTL on 6H (**Fig. 6D-E**). Most interestingly, the CS-SS QTL co-located with the QTL region identified for LS-IN on 6H (**Fig. 6C-E**), indicating that the 6H locus may regulate both phenotypes—CS-SS and LS-IN.

**Figure 6.**
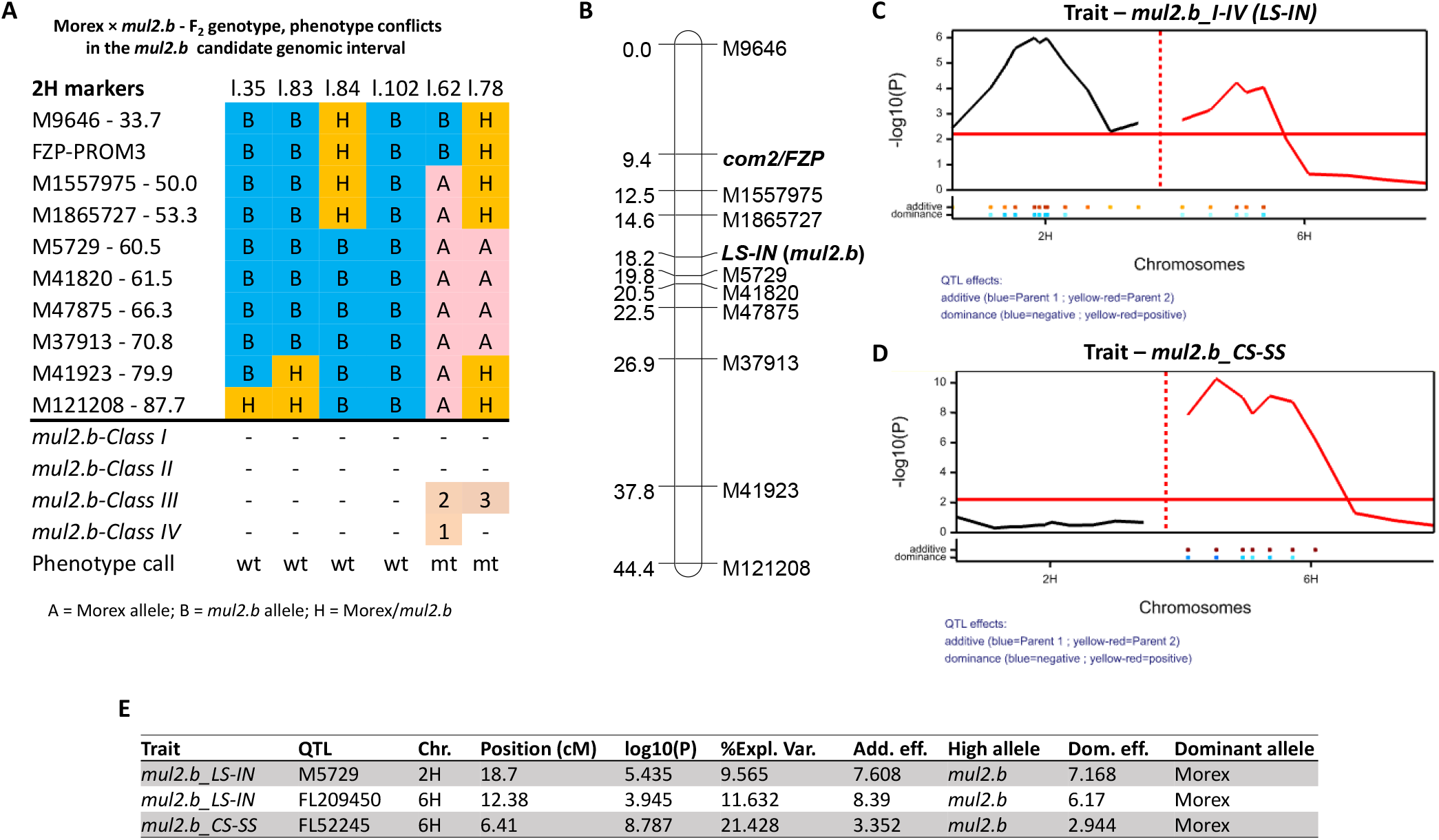
Linkage and QTL mapping of LS-IN and CS-SS phenotypes in Morex × *mul2.b* (POP 2015). (A) The upper panel shows the graphical genotypes of CAPS markers (derived from 2H LS-IN introgression identified from exome capture) that are not fitting with the LS-IN phenotype shown in the lower panel. The linkage map for rachilla elongation/additional floret phenotype after omitting the six lines with genotype-phenotype conflicts reported in figure 6A. (C) QTLs for rachilla elongation/additional floret on 2H, 6H chromosomes. (D) QTL for supernumerary spikelet phenotype on the 6H chromosome. (E) Summary of QTLs reported in figures 6C, and D.

### Whole-genome QTL analysis reveals more and novel genetic factors conditioning the LS-IN and CS-SS phenotypes

Since our partial-genome QTL analysis of LS-IN and CS-SS in the Morex × *mul2.b* population (POP 2015) indicated that these phenotypes are probably inherited in a quantitative fashion, we intended to perform whole-genome QTL analysis in the Morex × *mul2.b* F_2_ population. To this end, we screened another ~1,000 F_2s_ with markers M9646 and M41923 on 2H that are spaced apart by 37.8 cM (**Fig. 6B**). Between these two markers, we selected 177 recombinant individuals as an independent F_2_ population (POP 2016). We quantified their LS-IN and CS-SS phenotypes by counting the instances of LS-IN (cumulatively, number of *mul2-I, -II, -III*, and -*IV* classes) and CS-SS (extra central spikelets) per plant.

The LS-IN phenotype scored in POP 2016 fitted an incomplete dominant inheritance mode as observed in POP 2014-1 (χ^2^ = 0.092; p-value = 0.761). With respect to the CS-SS phenotype in POP 2016, 38 F_2s_ did not show a phenotype, whereas 138 F_2s_ carried the CS-SS phenotype. The majority of the F_2s_ harboring the CS-SS phenotype also had the LS-IN phenotype (104 F_2s_). The frequencies of phenotype distributions were also skewed for the LS-IN and CS-SS phenotypes, indicating that these traits were not strictly polygenic (**Fig. S5**). In spite of that, we proceeded analyzing LS-IN and CS-SS as quantitative traits, since we had observed two QTLs (on 2H and 6H chromosomes) for the LS-IN trait in the POP 2015. For the QTL analysis of F_2s_ in POP 2016, we normalized the phenotypic value of both traits (LS-IN and CS-SS) by estimating BLUEs to have a more meaningful view of the phenotypes on a per spike basis.

The 177 F_2s_ from POP 2016 were subjected to genotyping by sequencing (GBS) to generate SNP marker data. From our GBS analysis we got data for 1,573 high-quality SNP markers. Upon filtering for markers with >10% missing data or two or more linked markers without recombination across F_2s_, we mapped 605 markers onto 11 linkage groups. The barley chromosome 3H was split into three linkage groups, whereas 5H, 7H were split into two linkage groups due to lack of marker coverage mainly in the centromeric regions. Chromosomes 1H, 2H, 4H, and 6H formed unique linkage groups (**Supplementary table 3**). The genotypic and phenotypic data were used for QTL detection using Genstat 19.

Similar to the QTLs identified from POP 2015, we identified two major QTLs for LS-IN (cumulative *mul2.b-I, -II, -III, and -IV*) on 2H (termed *QMUL2.I-IV.ipk-2H*; -log10(P)–24.70; %PVE–32.19) and 6H (*QMUL2.I-IV.ipk-6H*; -log10(P)–10.93; %PVE–13.50) (**Fig. 7; Table 1**) in POP 2016. We also performed QTL analysis for the LS-IN component classes *mul2.b-I, -II, -III, and –IV* separately to further dissect the genetic basis of the individual phenotype classes. For the *mul2-I, -II*, and -*III* classes major QTLs were detected on 2H and 6H which co-located with *QMUL2.I-IV.ipk-2H* and *QMUL2.I-IV.ipk-6H* (**Fig. 7; Table 1**). For the weakest LS-IN component class - *mul2.b-IV*, only the 2H QTL co-located with both LS-IN cumulative and component QTLs on 2H while the 6H QTL was not detected for this class (**Fig. 7; Table 1**). Apart from the 2H and 6H QTLs for the *mul2.b-III* LS-IN component class, we identified two minor QTLs, *QMUL2.III.ipk-3H(1)* and *QMUL2.III.ipk-3H(2)* on 3H(1) and 3H(2) linkage groups explaining 8.43% and 5.65% of the observed phenotypic variation (**Fig. 7; Table 1**). Notably, in four out of five QTLs detected for the LS-IN trait the high phenotype value allele was derived from the *mul2.b* mutant parent; except for *QMUL2.III.ipk-3H(2)*, here the high-value allele was derived from Morex (**Fig. S6D**).

**Figure 7.**
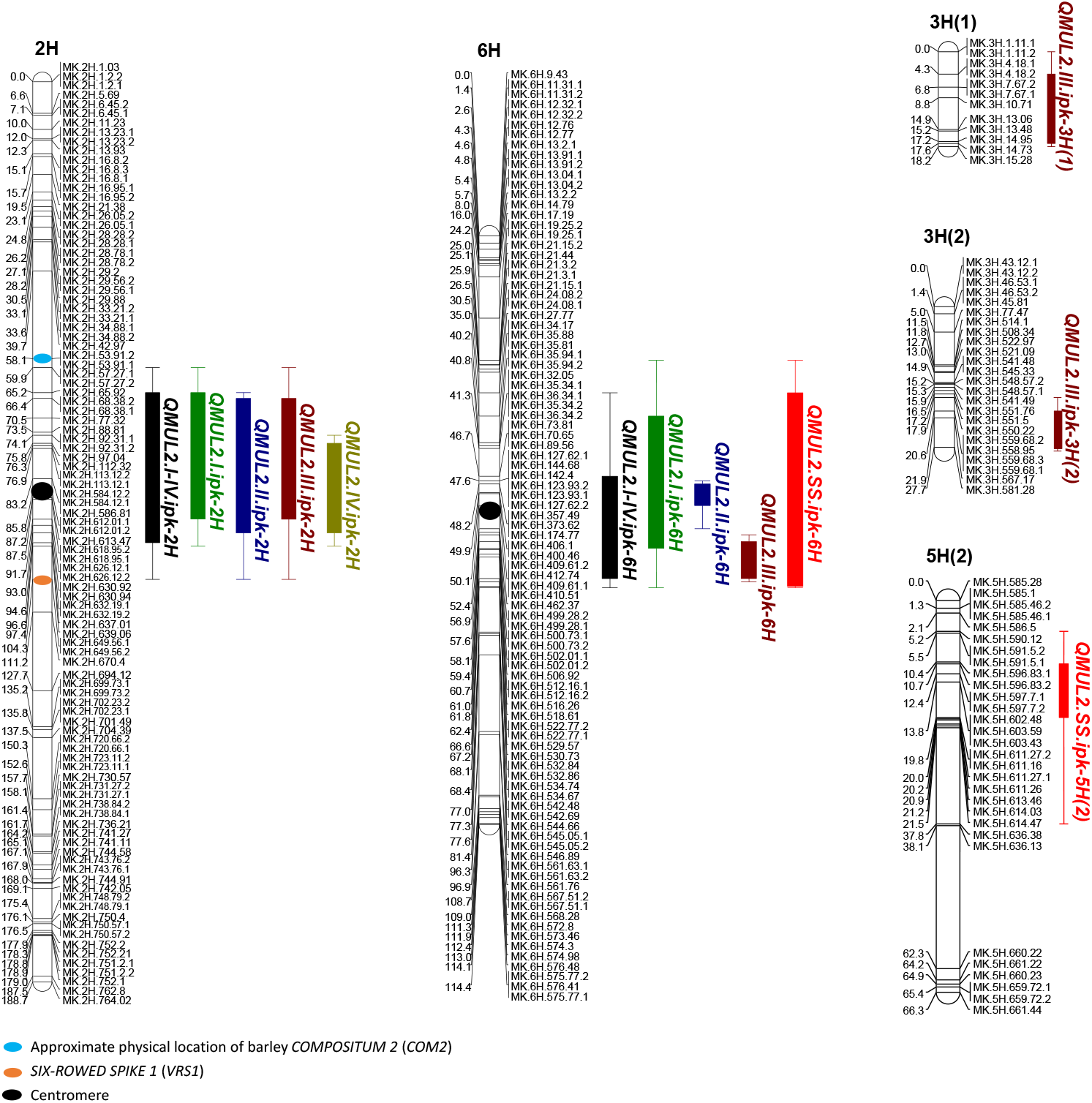
Identified QTLs for LS-IN and CS-SS phenotypes in the Morex × *mul2.b* (POP 2016)

**Table 1.**
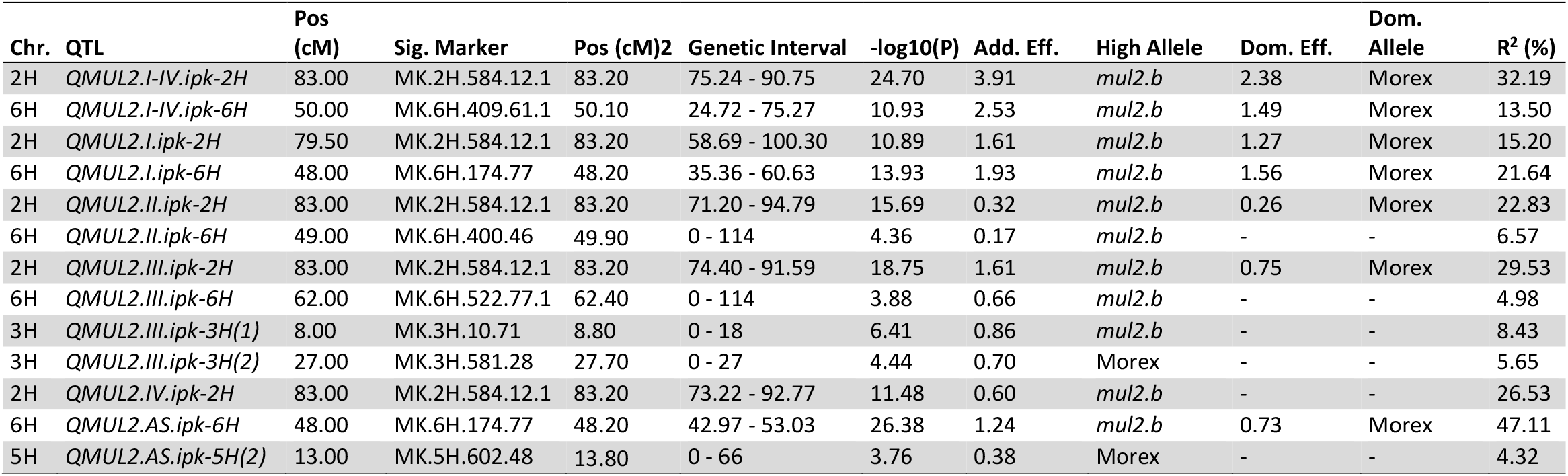
List of identified QTLs for LS-IN and CS-SS traits in the Morex × *mul2.b* (POP 2016)

For the CS-SS trait a single major QTL on 6H was identified (*QMUL2.SS.ipk-6H*; -log10(P)–26.38; %PVE – 47.11; **Fig. 7; Table 1**). Interestingly, *QMUL2.SS.ipk-6H* co-located with *QMUL2.I-IV.ipk-6H* and also with the 6H QTLs for LS-IN component classes, *mul2-I, -II*, and -*III* (**Fig.7**), indicating that a single genetic factor may underlie the LS-IN and CS-SS QTLs on 6H. We also found a minor QTL for the CS-SS trait on 5H(2) - *QMUL2.AS.ipk-5H(2)* explaining 4.32% of phenotypic variation (**Fig. 7; Table 1**). Interestingly, the high phenotype value allele for *QMUL2.AS.ipk-5H(2)* originated from the Morex parent (**Fig. S7B**). Thus, the Morex-derived high phenotype value alleles of minor QTLs, *QMUL2.III.ipk-3H(2)* for LS-IN and *QMUL2.AS.ipk-5H(2)* for CS-SS showed that independent genetic factors responsible for controlling these traits are present in both parents of the Morex × *mul2.b* cross.

### QTL allele effects for the LS-IN and CS-SS phenotypes

To estimate the phenotypic effect contributed by QTL alleles for the LS-IN trait, we evaluated combinations of Morex (“AA”), *mul2.b* (allele “*aa*”), and Morex/*mul2.b* (alleles *“A/a”*) alleles at *QMUL2.I-IV.ipk-2H* and *QMUL2.I-IV.ipk-6H* QTLs in F_2_. Interestingly, the strongest mutant LS-IN phenotype (cumulative *mul2.b-I, -II, -III, and -IV*) was manifested only in F_2s_ with mutant “*aa*” alleles present in both QTL loci (**Fig. S8A**). The LS-IN phenotypic effect got drastically reduced in F_2s_ with alleles “*aa*” and “*AA*” at 2H and 6H QTLs, respectively (**Fig. S8A**). Also heterozygosity at both QTL loci in combination with the respective mutant “*aa*” allele [e.g. *aa*(2H)/*Aa*(6H); or *Aa*(2H)/*aa*(6H)] enhanced LS-IN phenotype in comparison with the wild type allele combination, [(e.g. *aa*(2H)/*AA*(6H) or *AA*(2H)/*aa*(6H)] indicating that *QMUL2.I-IV.ipk-6H* enhanced *QMUL2.I-IV.ipk-2H* effect in an additive manner (**Fig. S8A**). The single major QTL for the CS-SS trait (*QMUL2.SS.ipk-6H*), however, was not influenced by the alleles at *QMUL2.I-IV.ipk-2H*, indicating that the CS-SS trait is majorly controlled by 6H QTL **(Fig. S8B**). Similar phenotypic effects for LS-IN and CS-SS traits were also observed in F_2:3_ generation (**Fig. S9**), indicating the stable inheritance of these phenotypes.

The LS-IN component trait *mul2.b-III* had two major QTLs, one each on 2H (*QMUL2.III.ipk-2H*) and 6H (*QMUL2.III.ipk-6H*) but also two minor QTLs on 3H (*QMUL2.III.ipk-3H(1), QMUL2.III.ipk-3H(2)*). Interestingly, the phenotypic effect was highly enhanced in F_2s_ with favorable QTL alleles at these four loci, i.e. *mul2.b* allele at 2H, 6H, 3H(1) QTLs, and Morex allele at 3H(2) QTL, indicating the additivity of these loci (**Fig. S6**). Also for the CS-SS trait, a similar additive phenotype was observed in F_2_ individuals carrying high phenotype value alleles at *QMUL2.SS.ipk-6H* (*mul2.b* alleles) and *QMUL2.SS.ipk-5H(2)* (Morex alleles) loci (**Fig. S7**.)

## Discussion

### Mutations in multiflorus.2b induce two ancestral phenotypes – LS-IN and CS-SS

The spikes of the *Triticeae* tribe are proposed to be derivatives of ancestral compound spike/panicle inflorescences obtained through an evolutionary series of inflorescence branch complexity reduction (Koppolu and Schnurbusch 2019; Perreta et al. 2009; Vegetti 1995). Within Triticeae species, spike inflorescences show structural differences in terms of determinacy at the levels of the spike (presence or absence of terminal spikelet) and spikelet (uni- or multi-floreted spikelet) (Sakuma et al. 2011). The uni-floreted condition is manifested due to suppression of rachilla elongation after the initiation of the first floret, whereas in multi-floreted spikelets rachilla elongation continues to generate more florets. Two peculiar Triticeae species with contrasting determinacy features at both levels include barley (determinate spikelet, indeterminate spike) and wheat (determinate spike, indeterminate spikelet). It was proposed that terminal spikelet identity (determinate spike) and multi-floreted conditions (indeterminate spikelet) of wheat are ancestral, whereas indeterminate spike and uni-floreted (determinate spikelet) conditions are derived features in barley (Clayton 1990; Zhong et al. 2021). The mutants of barley *mul2.b* lost the spikelet determinacy producing ancestral multi-floreted condition similar to wheat spikelets, but the derived spike indeterminacy character is retained. Interestingly the multi-floreted condition was restricted to LS of *mul2.b* (LS-IN) with CSs retaining the determinate character. Such determinacy differences between CS and adjacent LSs could be probably due to separate genetic factors controlling this condition in CS and LS differently.

Notably, mutants of *mul2.b* also produce supernumerary spikelets (SS) in the CS (CS-SS). The SS feature appears to be analogous with the paired spikelet phenotype of wheat and the spikelet pair phenotype characteristic to maize, sorghum, and other *Andropogoneae* species (Boden et al. 2015). Also, species belonging to the Triticeae genera *Crithops* and *Taeniatherum* bear two spikelets at each rachis node on the spike (Frederiksen and Seberg 1992), probably indicating that the mechanism regulating SS phenotype is an ancestral feature retained in some of the *Triticeae* species. However, the CS-SS phenotype in *mul2.b* and *flo.a* could be potentially a revertant ancestral phenotype due to mutations in respective loci.

### *LS-IN and CS-SS are quantitatively inherited in the Morex × mul2.b* cross

Based on our in-depth phenotypic analysis we showed that the *mul2.b* phenotype is a combination of rachilla elongation in LSs (LS-IN) and supernumerary spikelet formation in CSs (CS-SS). The previous genetic and phenotypic analyses involving *mul2.b* apparently did not consider the CS-SS phenotype in central spikelets. Also, the LS-IN phenotype was not evaluated in sufficient detail probably owing to difficulties in visualizing the rachilla elongation in lateral spikelets. This might have potentially lead to the classification of *mul2.b* as a monogenic recessive locus (Walker et al. 1963). Our QTL analysis based on comprehensive LS-IN and CS-SS phenotypes revealed two major QTLs for LS-IN (*QMUL2.I-IV.ipk-2H, QMUL2.I-IV.ipk-6H*) and one major QTL for the CS-SS phenotype (*QMUL2.SS.ipk-6H*) (**Fig. 7; Table 1**). Also, two minor QTLs for LS-IN [*QMUL2.III.ipk-3H(1), QMUL2.III.ipk-3H(2)*] and one for CS-SS phenotype [*QMUL2.SS.ipk-5H(2)*] were identified indicating the quantitative nature of the *mul2.b* phenotypes (**Fig. 7; Table 1**). It should be noted that the cv. Morex (wt parent) occasionally showed mild and infrequent rachilla elongation in LSs as well as supernumerary spikelet formation in CS (data not shown). Consistent with this, we observed for each trait one minor QTL, *QMUL2.III.ipk-3H(2)* and *QMUL2.SS.ipk-5H(2)*, where the phenotypic high-value allele was derived from Morex (**Fig. 7; Table 1**). Previous genetic analyses in wheat involving the paired spikelet phenotype (analogous to *mul2.b* CS-SS) also reported quantitative variation for this trait (Boden et al. 2015; Dobrovolskaya et al. 2015; Echeverry-Solarte et al. 2014).

Our genetic analysis of LS-IN and CS-SS phenotypes revealed co-locating QTLs on 6H (*QMUL2.I-IV.ipk-6H, QMUL2.SS.ipk-6H*; **Fig. 7**), indicating that these phenotypes are potentially regulated by a single locus on 6H. On top of that, the phenotypic effect of LS-IN QTL on 2H is significantly enhanced by the 6H QTL alleles in an additive manner, without which, the LS-IN phenotypic was drastically reduced (**Fig. S7**). Interestingly, the Bowman near-isogenic line of *mul2.b* (BM-NIL(*mul2.b*); GSHO 2089, BW607) showed a mild LS-IN phenotype, whereas the CS-SS phenotype is completely missing in BM-NIL(*mul2.b*) (**Fig. S10**). Such a mild LS-IN phenotype and lack of CS-SS phenotype in BM-NIL(*mul2.b*) could be probably due to the lack of *mul2.b* 6H introgression from the original mutant that enhances the phenotypic effect of 2H QTL. For the LS-IN phenotype component phenotype class *mul2.b-III* we also identified two minor QTLs on 3H. Thus, our precise phenotype quantification data guided us to genetically dissect genomic regions responsible for various LS-IN phenotypes.

### Conclusion

Yield in small grain cereals such as barley and wheat is mainly contributed by the number of grains produced per unit area (Ferrante et al. 2017; García et al. 2015; Sakuma and Schnurbusch 2020; Serrago et al. 2013). The grain number can be improved by increasing spikelet number per spike, floret number per spikelet and also enhancing floret fertility within spikelets. Here we characterized a barley mutant, *mul2.b*, that showed multi-floreted spikelets as well as more spikelets per spike (supernumerary spikelet formation). Through our genetic analysis, we identified component QTLs conditioning these two trait phenotypes. Further, we laid the foundation for identifying and characterizing the genetic factors underlying these phenotypes by QTL cloning in the future.

## Acknowledgments

The authors thank Udda Lundqvist, Jerome Franckowiak, and Thirulogachandar Venkatasubbu for the insightful discussions on the *multiflorus2.b* mutant. We thank Harold Bockelman (USDA-ARS, USA) for providing the *multiflorus2.b* and Montcalm germplasm. We thank Roopkamal Kaur for help with the data analysis. We thank Corinna Trautewig, Mechthild Pürschel, Angelika Pueschel, Kerstin Wolf, Susanne König and Manuela Knauft for excellent technical assistance. We thank Anne Fiebig for GBS and exome capture sequence data submission. We are also thankful to Heike Müller and Andreas Bähring for supporting graphical artwork.

## Funding

This study received financial support from the Chinese Scholarship Council (CSC) to G.J. The Schnurbusch laboratory was supported by the HEISENBERG Program of the German Research Foundation (DFG) (grant nos. SCHN 768/8-1 and SCHN 768/15-1), the European Research Council (ERC) (grant agreement no. 681686 “LUSH SPIKE”), ERC-2015-CoG, and IPK core budget.

## Conflicts of interest

Authors declare no conflicts of interest.

## Availability of data and material

The sequence data generated in this study can be accessed from European Nucleotide Achive (ENA) under the accession numbers PRJEB44305 (exome capture) and PRJEB44306 (GBS).

## Authors’ contributions

R.K. conceptualized the study together with T.S.; T.S. and R.K. jointly supervised the study. R.K. generated mapping populations and established the *mul2.b* phenotyping methodology. R.K. and G.J. conducted phenotypic and genetic analyses. Q.H.M. generated BLUE scores for POP 2016. S.G.M. conducted GBS data analysis for POP 2016. T.R. conducted scanning electron microscopy of immature barley spikes. A.H. and N.S. provided expertise and resources for exome capture and GBS experiments. M.M. conducted exome capture data analysis for POP 2014. R.K. drafted the MS with the help from T.S. All authors read and approved the MS.

## Supplementary figures and tables

**Supplementary figure 1.**
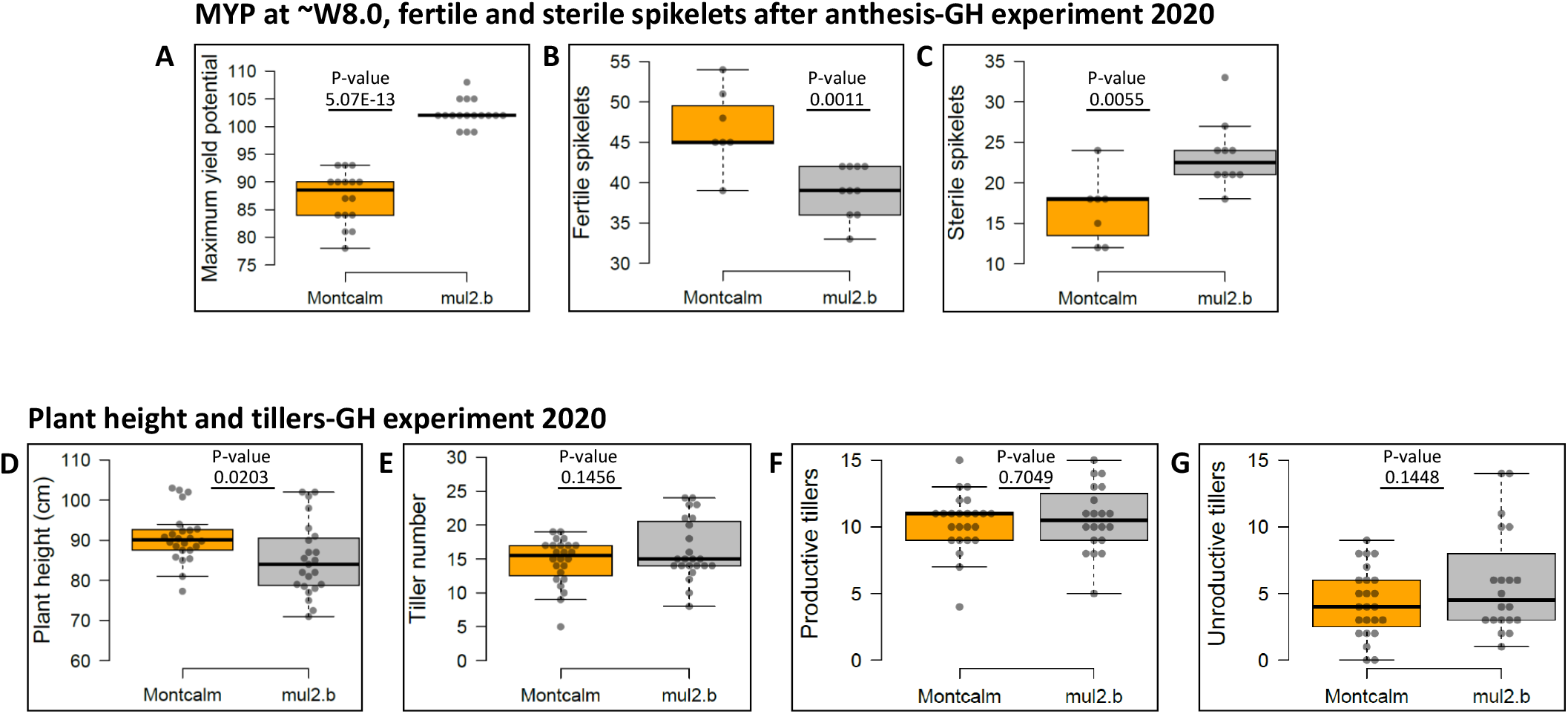
Spike and biomass-related traits – Montcalm vs. *mul2.b*. (A) Maximum yield potential of Montcalm and *mul2.b* measured before anthesis. (B-C) Fertile grain-bearing spikelets (B) and sterile spikelets (C) at spike maturity. (D-G) Biomass-related traits such as plant height (D) total tillers (E), productive (F), and unproductive (G) tillers. P-values were calculated based on *Student’s t-test* (two-tailed).

**Supplementary figure 2.**
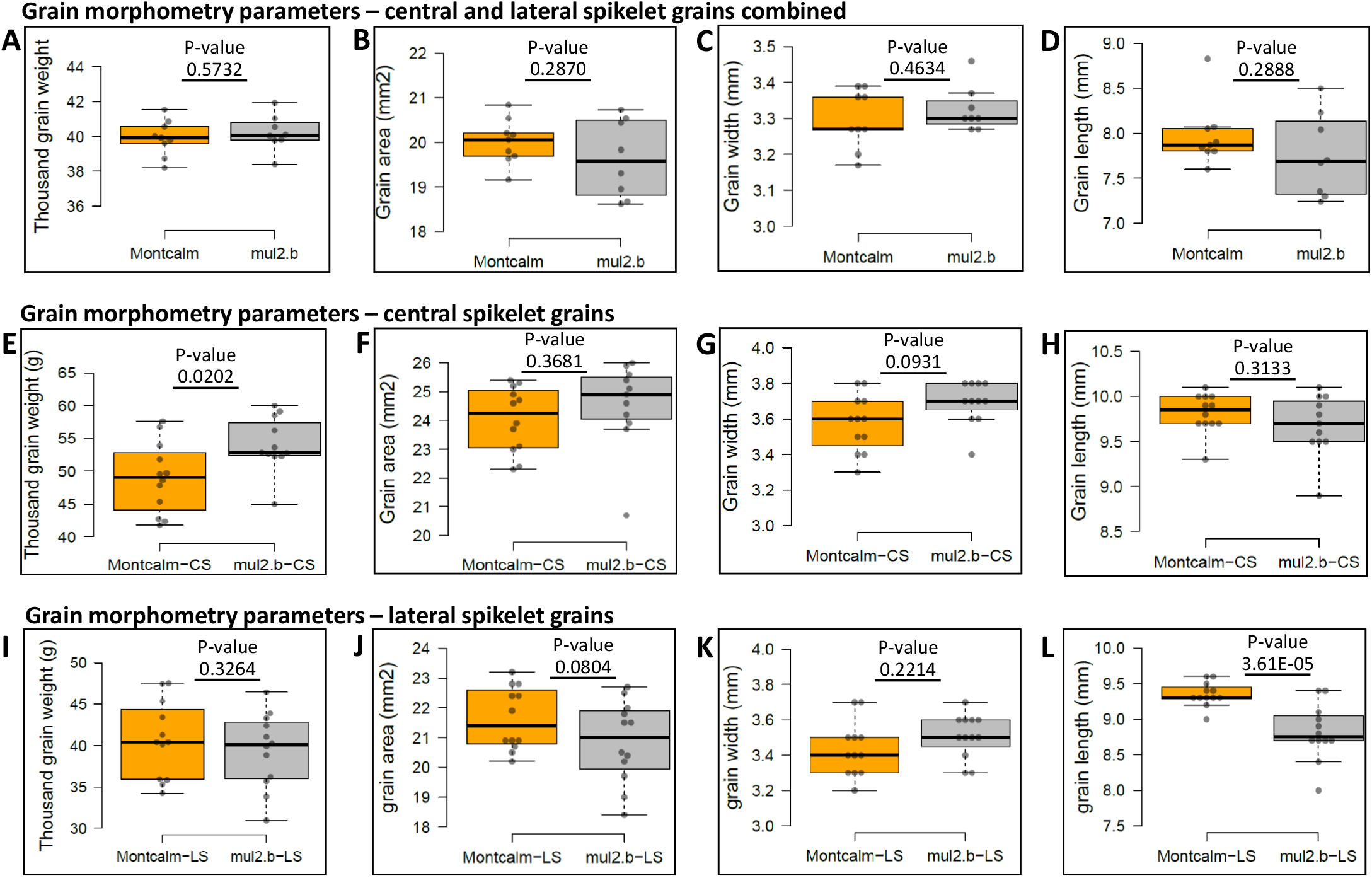
Grain morphometry parameters – Montcalm vs. *mul2.b*. (A-D) Thousand-grain weight (A), grain area (B), width (C), and length (D) of Montcalm and *mul2.b* measured from central and lateral spikelet grains together. (E-H) Thousand-grain weight (E), grain area (F), width (G), and length (H) of Montcalm and *mul2.b* measured from central spikelet grains. (I-L) Thousand-grain weight (I), grain area (J), width (K), and length (L) of Montcalm and *mul2.b* measured from lateral spikelet grains. P-values were calculated based on *Student’s t-test* (two-tailed).

**Supplementary figure 3.**
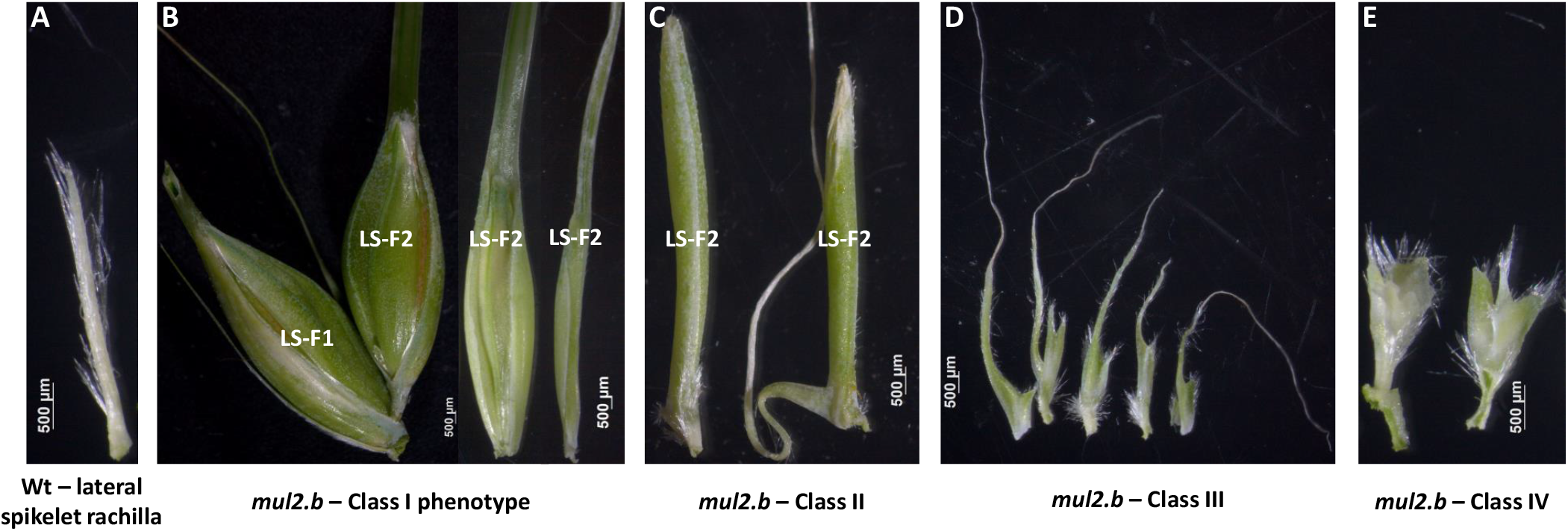
*mul2.b* phenotype classification for lateral spikelet indeterminacy. (A) Wild-type rachilla of Montcalm. (B-C) class I (B) and class II (C) additional florets excised from the lateral spikelets of *mul2.b*. (D-E) Malformed rachillae of Class III (D) and class IV (E) excised from lateral spikelets.

**Supplementary figure 4.**
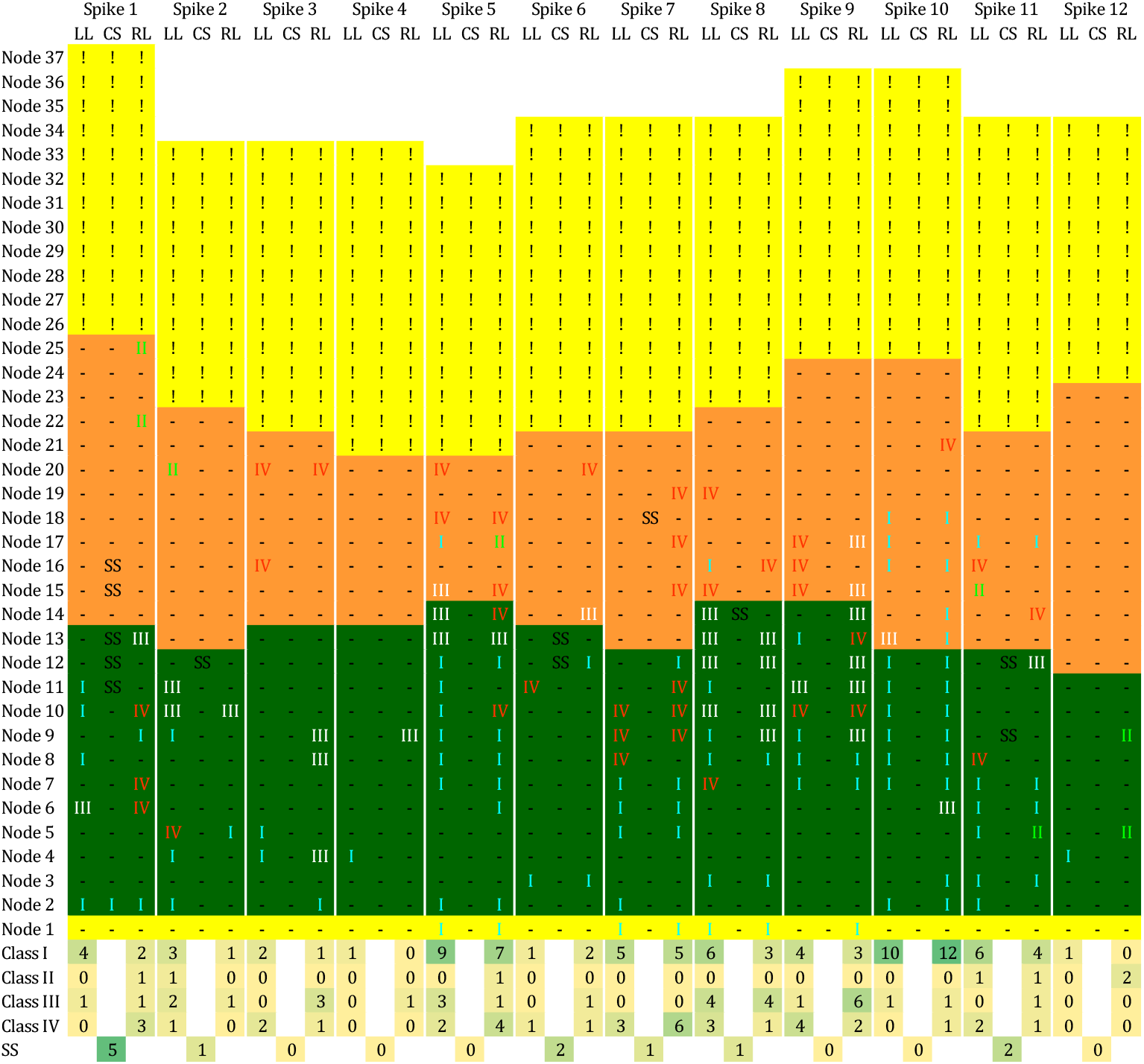
Distribution of different classes of lateral spikelet indeterminacy and supernumerary spikelet phenotypes in spikes of the *mul2.b* mutant. The yellow-colored cells at spikelet node 1 and in the upper spikelet nodes indicate the aborted spikelets at the base and top of the spike, respectively after anthesis. Cells highlighted in green indicate the fertile spikelets that potentially bear a grain. The cells in orange indicate the unfilled spikelets (spikelets formed but did not grow further). Classes, I, II, III, and IV indicate respective phenotypes for rachilla indeterminacy or additional floret formation in lateral spikelets, whereas SS is supernumerary spikelet phenotype in central spikelets. The notation “!” in the upper aborted spikelets (Yellow) indicates the absence of phenotypic information for lateral spikelet indeterminacy/additional spikelet and supernumerary spikelet phenotypes. CS – Central Spikelet; LL – Left Lateral spikelet; RL – Right Lateral spikelet; SS – Supernumerary spikelet.

**Supplementary figure 5.**
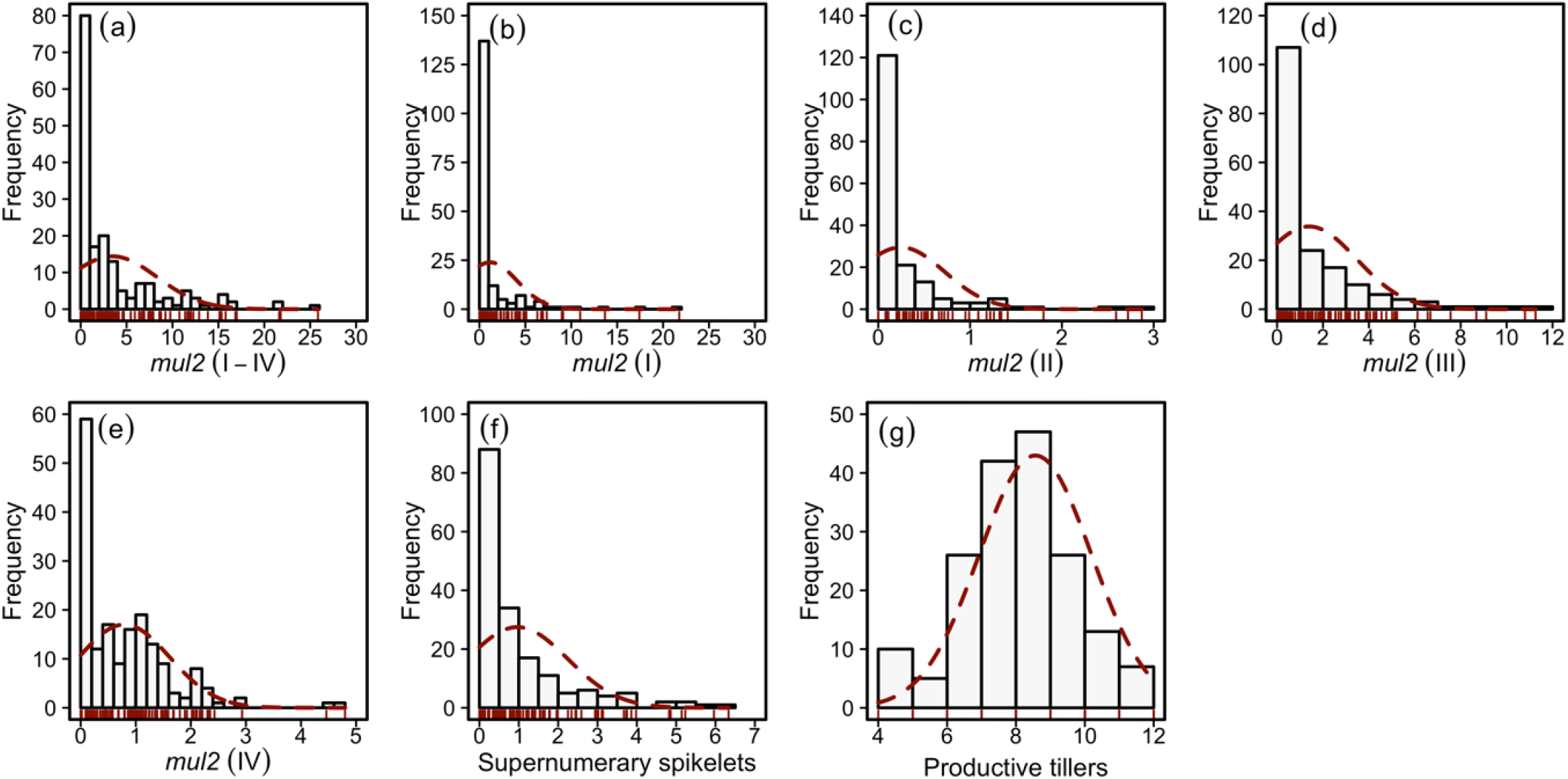
Frequency distribution of LS-IN and CS-SS phenotypes in Morex × *mul2.b* (POP 2016)

**Supplementary figure 6.**
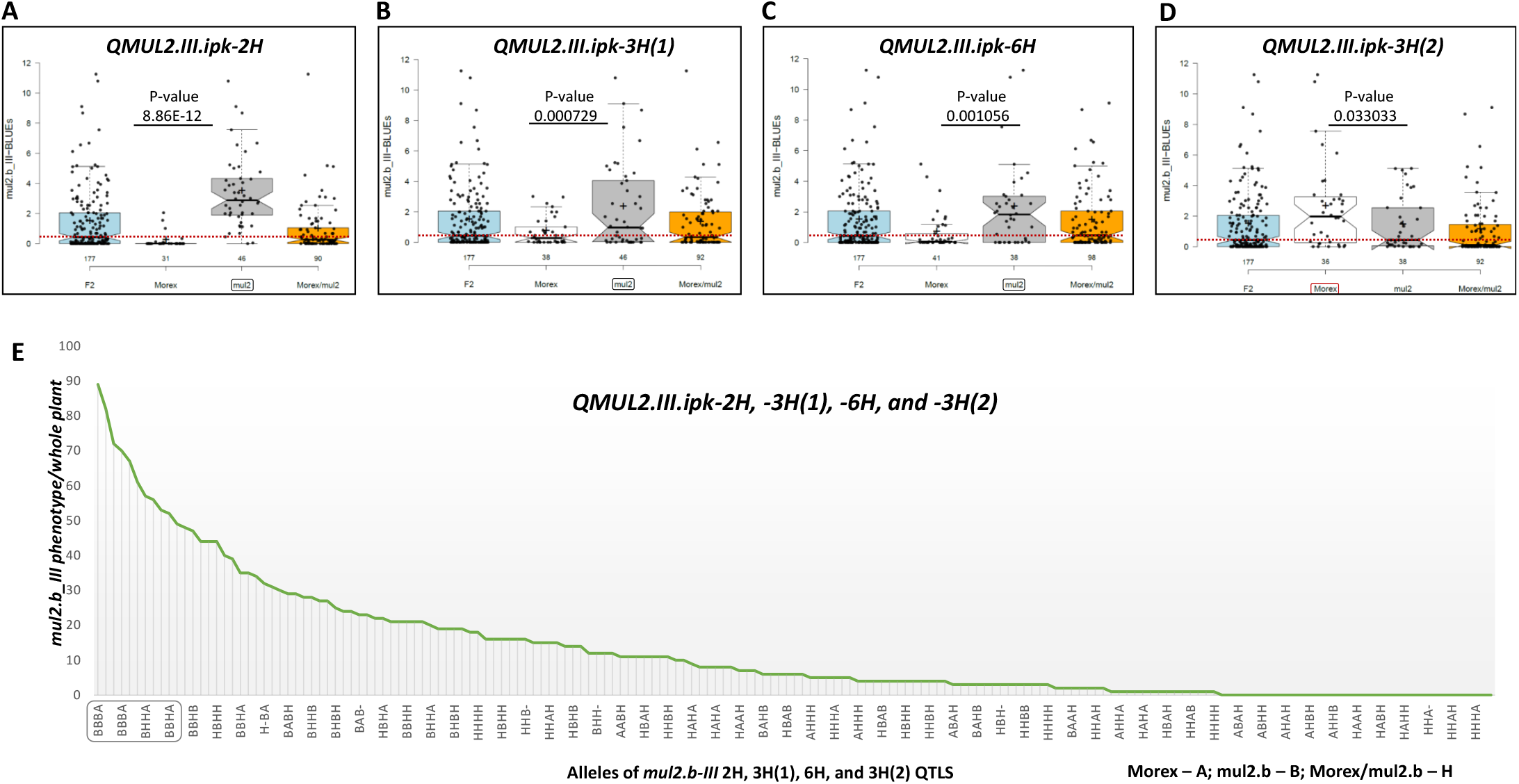
Phenotypic effect of LS-IN trait at individual QTLs *QMUL2.III.ipk-2H* (A) *QMUL2.III.ipk-3H(1*) (B) *QMUL2.III.ipk-6H* (C) and *QMUL2.III.ipk-3H(2)* (D). (E) The cumulative phenotypic effect from the four QTLs for class III LS-IN phenotype. Phenotype frequencies shown in the box plots are derived from the genotypes of markers with the highest −log10 (p-value) and respective trait phenotype values.

**Supplementary figure 7.**
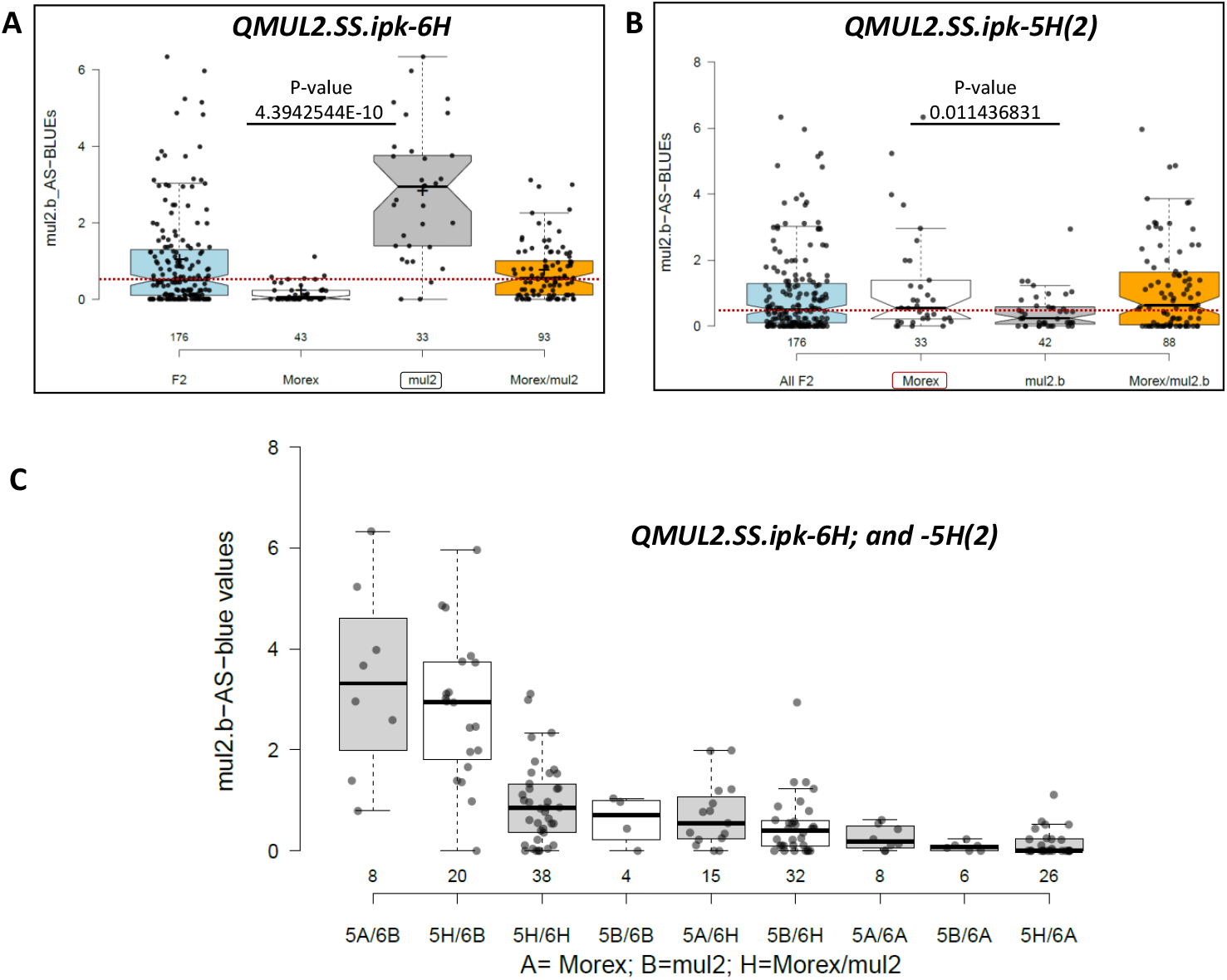
Phenotypic effect of CS-SS trait at individual QTLs *QMUL2.SS.ipk-6H* (A) and *QMUL2.SS.ipk-5H(2)* (B). (C)The cumulative phenotypic effect from both QTLs. Phenotype frequencies shown in the box plots are derived from the genotypes of markers with the highest −log10 (p-value) and respective trait phenotype values.

**Supplementary figure 8.**
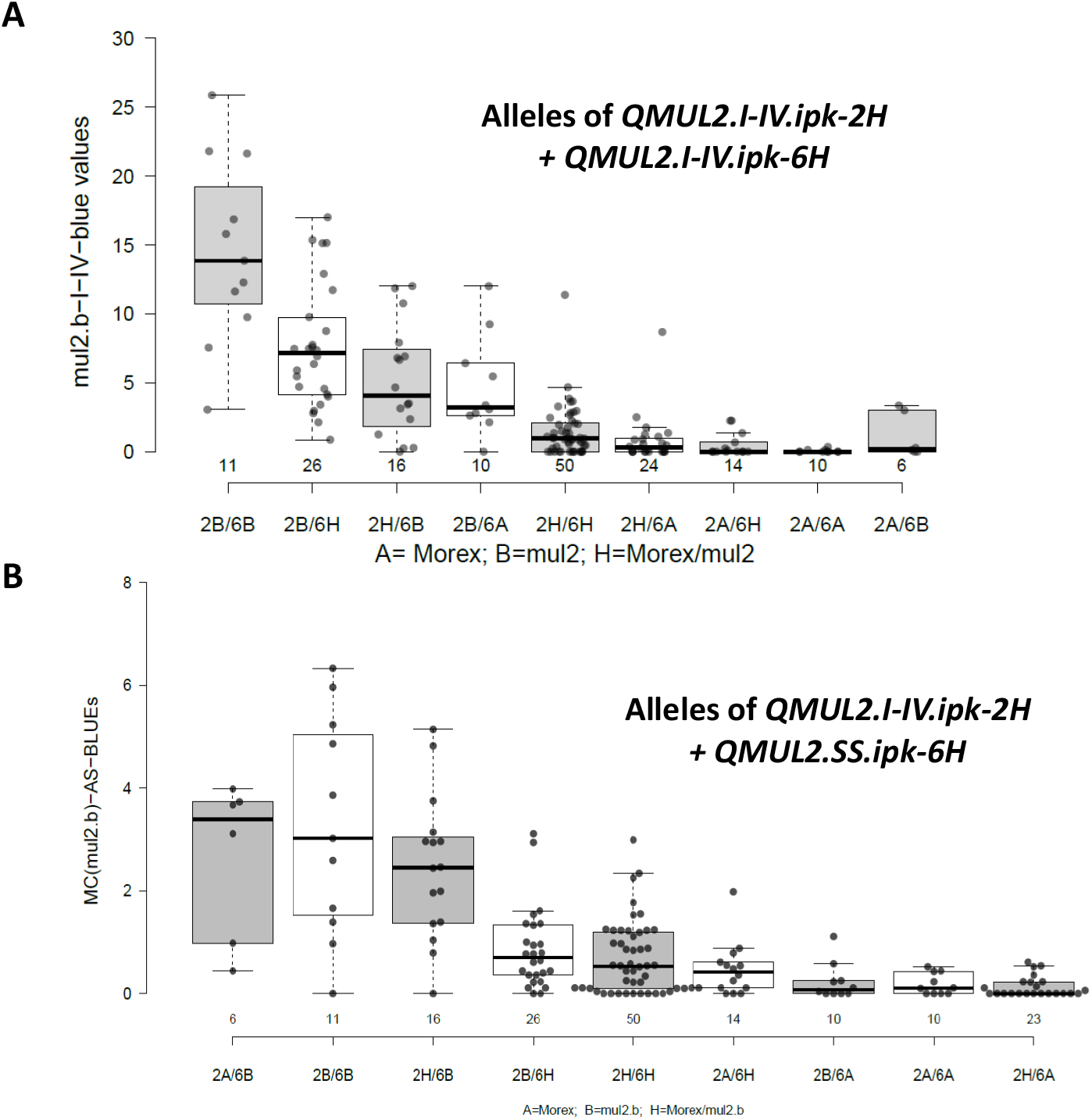
Phenotypic effect of LS-IN and CS-SS QTLs in Morex × *mul2.b*-F_2_. (A) Cumulative phenotypic effect of *QMUL2.I-IV.ipk-2H* and *QMUL2.I-IV.ipk-6H* QTL alleles (B) Cumulative phenotypic effect *QMUL2.I-IV.ipk-2H* and *QMUL2.SS.ipk-6H* QTL alleles. Phenotype frequencies shown in the box plots are derived from the genotypes of markers with the highest −log10 (p-value) and respective trait phenotype values.

**Supplementary figure 9.**
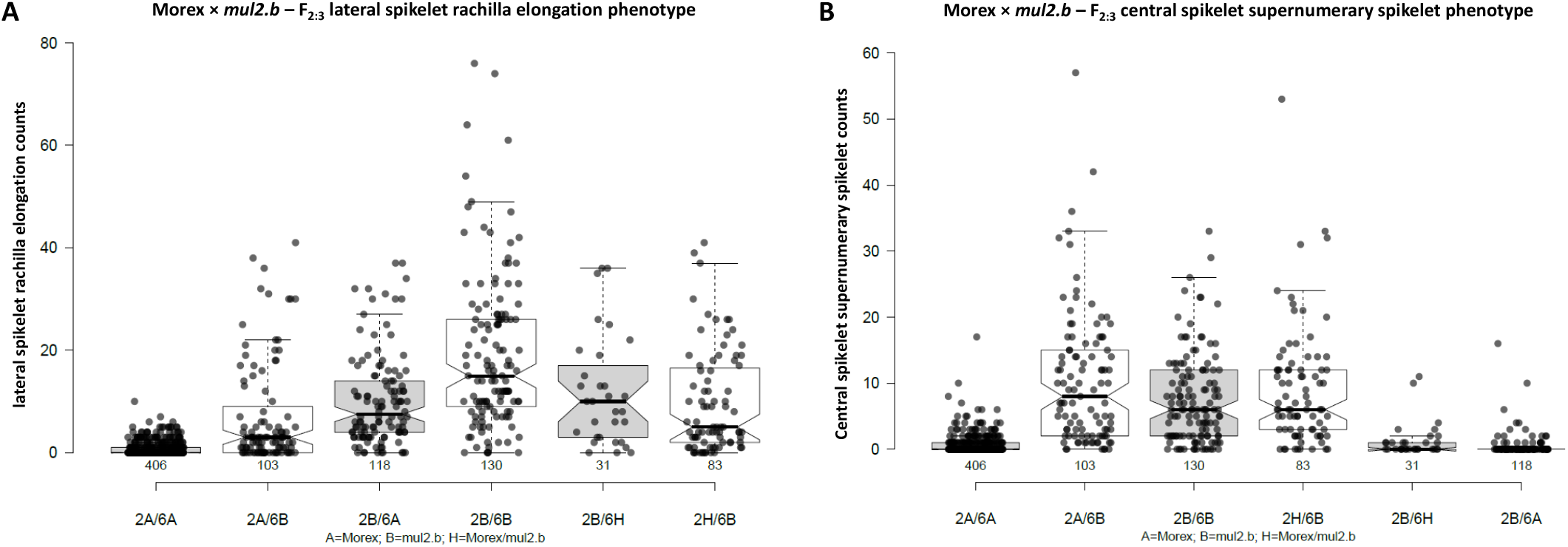
Phenotypic effect of LS-IN (A) and CS-SS traits (B) in F_2:3_ progenies of Morex × *mul2.b*. The markers explaining the highest phenotypic value for the respective traits at 2H and 6H QTL were considered for the analysis.

**Supplementary figure 10.**
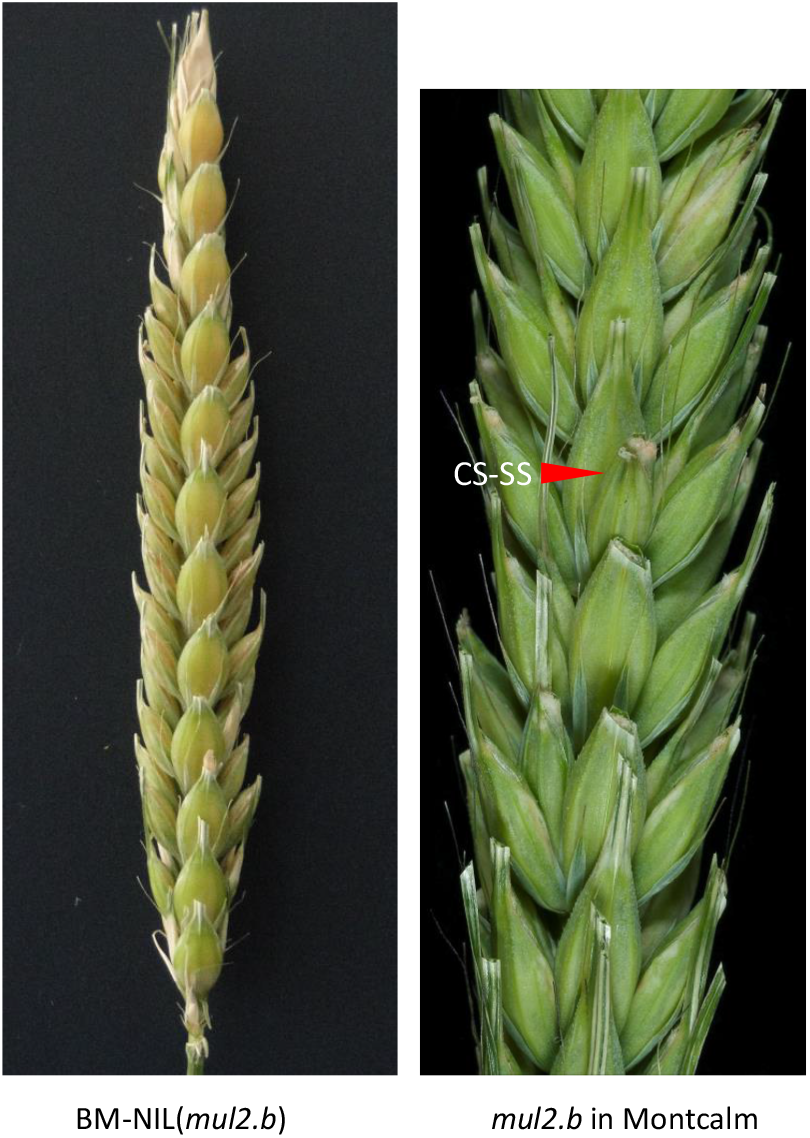
CS-SS phenotype comparison beteen BM-NIL(*mul2.b*) and *mul2.b* in wildtype progenitor Montcalm.

**Supplementary Table 1. Sequences of primers used for marker mapping in Morex × *mul2.b***

**Supplementary Table 2. Segregation pattern of LS-IN phenotype in Morex × *mul2.b* populations.**

**Supplementary Table 3. Genetic linkage map features-Morex × *MC*(*mul2.b*) – POP 2016.**

## Notes

### Competing Interest Statement

The authors have declared no competing interest.

